# Neuronal DAMPs exacerbate neurodegeneration via astrocytic RIPK3 signaling

**DOI:** 10.1101/2023.07.21.550097

**Authors:** Nydia P. Chang, Evan M. DaPrano, Wesley R. Evans, Marialaina Nissenbaum, Micheal McCourt, Diego Alzate, Marissa Lindman, Tsui-Wen Chou, Colm Atkins, Alexander W. Kusnecov, Rafiq Huda, Brian P. Daniels

## Abstract

Astrocyte activation is a common feature of neurodegenerative diseases. However, the ways in which dying neurons influence the activity of astrocytes is poorly understood. RIPK3 signaling has recently been described as a key regulator of neuroinflammation, but whether this kinase mediates astrocytic responsiveness to neuronal death has not yet been studied. Here, we used the MPTP model of Parkinson’s disease to show that activation of astrocytic RIPK3 drives dopaminergic cell death and axon damage. Transcriptomic profiling revealed that astrocytic RIPK3 promoted gene expression associated with neuroinflammation and movement disorders, and this coincided with significant engagement of DAMP signaling. Using human cell culture systems, we show that factors released from dying neurons signal through RAGE to induce RIPK3-dependent astrocyte activation. These findings highlight a mechanism of neuron-glia crosstalk in which neuronal death perpetuates further neurodegeneration by engaging inflammatory astrocyte activation via RIPK3.

## Introduction

Recent work has identified a central role for neuroinflammation in the pathogenesis of neurological disease, including major neurodegenerative disorders such as Alzheimer’s and Parkinson’s disease^1,2^. Although glial cells are critical regulators of neuroinflammation, activated glia serve complex roles during disease, including both protective and pathogenic functions^3^. Among glial cells, astrocytes are the most abundant cell type in the central nervous system (CNS), where they support homeostasis via wide-ranging effects on neurotransmission, neurovascular function, and metabolism^4^. However, following an inflammatory insult, astrocytes can enter “reactive” states associated with disease pathogenesis^5^. While astrocyte activation is likely highly plastic and context-dependent, it is now widely accepted that astrocytes can take on inflammatory transcriptional states during disease that are associated with the conferral of neurotoxic activity and suppression of normal homeostatic functions^6^. Despite this understanding, the molecular mechanisms that govern astrocyte reactivity during neurodegenerative disease, and particularly those factors that most directly exacerbate disease progression, remain poorly understood^7^.

We and others have recently identified receptor-interacting serine/threonine protein kinase-3 (RIPK3) as a key regulator of inflammation in the CNS^8-10^. RIPK3 signaling is canonically associated with necroptotic cell death, which is induced via the activation of mixed lineage kinase domain-like protein (MLKL)^11^. While RIPK3-dependent necroptosis has been implicated in the pathogenesis of several neurological disorders, RIPK3 also appears to promote neuroinflammatory processes via necroptosis-independent mechanisms, including the coordination of inflammatory transcription in multiple CNS cell types^12-18^. While necroptosis-independent roles for RIPK3 signaling in astrocytes have not been thoroughly studied, we have previously shown that pathogenic α-synuclein fibrils activate RIPK3 signaling in human midbrain astrocyte cultures, resulting in NF-κB-mediated transcriptional activation without inducing astrocytic necroptosis^14^. However, whether RIPK3 controls astrocyte transcriptional activation and function in models of neurodegenerative disease *in vivo* is unknown.

The importance of neuron-glia communication during CNS disease states has also gained significant recognition in recent work^19-22^. A particularly important goal in this area is defining the stimuli that induce inflammatory signaling in the “sterile” setting of neurodegeneration. One potential stimulus underlying inflammatory astrocyte activation during neurodegeneration are factors derived from dead and dying neurons, themselves. These factors include damage-associated molecular patterns (DAMPs), molecules released from damaged cells that serve as endogenous danger signals that elicit potent innate immune activation in neighboring cells^23,24^. DAMP release has been associated with numerous inflammatory diseases, including neurodegenerative disorders^25-28^. However, whether and how neuron-derived DAMPs impact astrocyte function during neurodegenerative disease has not been thoroughly studied to date.

Here, we define a new role for RIPK3 signaling in mediating astrocyte activation downstream of neuronal DAMP release. We utilize the 1-methyl-4-phenyl-1,2,3,6-tetrahydropyridine (MPTP) model of Parkinson’s disease, in which cell death can be selectively induced in dopaminergic neurons *in vivo*, to show that induction of neuronal cell death results in RIPK3-dependent astrocyte activation, which in turn exacerbates ongoing neurodegeneration. Transcriptional profiling revealed a robust RIPK3-dependent inflammatory signature in astrocytes exposed to dying neuron-derived factors, and this occurred independently of astrocytic MLKL. Mechanistically, we show that factors released from dying dopaminergic neurons activate receptor for advanced glycation endproducts (RAGE) on midbrain astrocytes. RAGE signaling, in turn, was required for RIPK3 activation, inflammatory transcription, and the conferral of neurotoxic activity in midbrain astrocytes following exposure to neuronal DAMPs. Our findings suggest a feed-forward mechanism that perpetuates neurodegeneration via the DAMP-dependent activation of RIPK3-dependent inflammation and neurotoxicity in astrocytes. These results highlight an important mechanism of neuron-glia crosstalk, with implications for the prevention and treatment of neurodegenerative disease.

## Results

### Astrocytic RIPK3 signaling promotes pathogenesis in the MPTP model of Parkinson’s disease

To examine the impact of astrocytic RIPK3 signaling in response to neuronal cell death, we subjected mice with astrocyte-specific deletion of *Ripk3* (*Ripk3*^fl/fl^ *Aldh1l1*^Cre+^) and littermate controls to treatment with MPTP, a neurotoxin that selectively induces death in dopaminergic neurons^29,30^. MPTP administration resulted in significant loss of tyrosine hydroxylase (TH) immunoreactivity in the substantia nigra pars compacta (SNpc) of control animals, consistent with the depletion of dopaminergic neurons in this region (Figure 1A-B). Strikingly, however, *Ripk3*^fl/fl^ *Aldh1l1*^Cre+^ mice exhibited greatly reduced dopaminergic neuron loss following MPTP treatment, suggesting a role for astrocytic RIPK3 in exacerbating neuronal death in this model. We also observed a significant loss of TH^+^ dopaminergic axons in the striatum of control animals (Figure 1C-D), along with increased frequencies of TH^+^ axons immunoreactive for SMI32, a marker of axonal degeneration^31-33^ (Figure 1E). This phenotype was also greatly ameliorated in *Ripk3*^fl/fl^ *Aldh1l1*^Cre+^ mice. To test whether these differences in dopaminergic neuron loss were associated with differences in motor function, we next subjected mice to the vertical grid maze, a motor task previously shown to be sensitive to perturbations of dopaminergic circuits^34,35^. Strikingly, MPTP-treated control mice exhibited significantly impaired performance in the vertical grid maze (Figure 1F-G), while mice lacking astrocytic *Ripk3* did not. Improvements in dopaminergic neuron loss and motor performance in *Ripk3*^fl/fl^ *Aldh1l1*^Cre+^ mice were not due to differential metabolism of MPTP compared to Cre- littermates, as we observed indistinguishable levels of the toxic metabolite of MPTP (MPP^+^) in midbrain homogenates derived from animals of both genotypes (Supplemental Figure 1). Together, these data suggest that astrocytic RIPK3 signaling exacerbates neuronal cell death following a neurotoxic insult.

**Figure 1.**
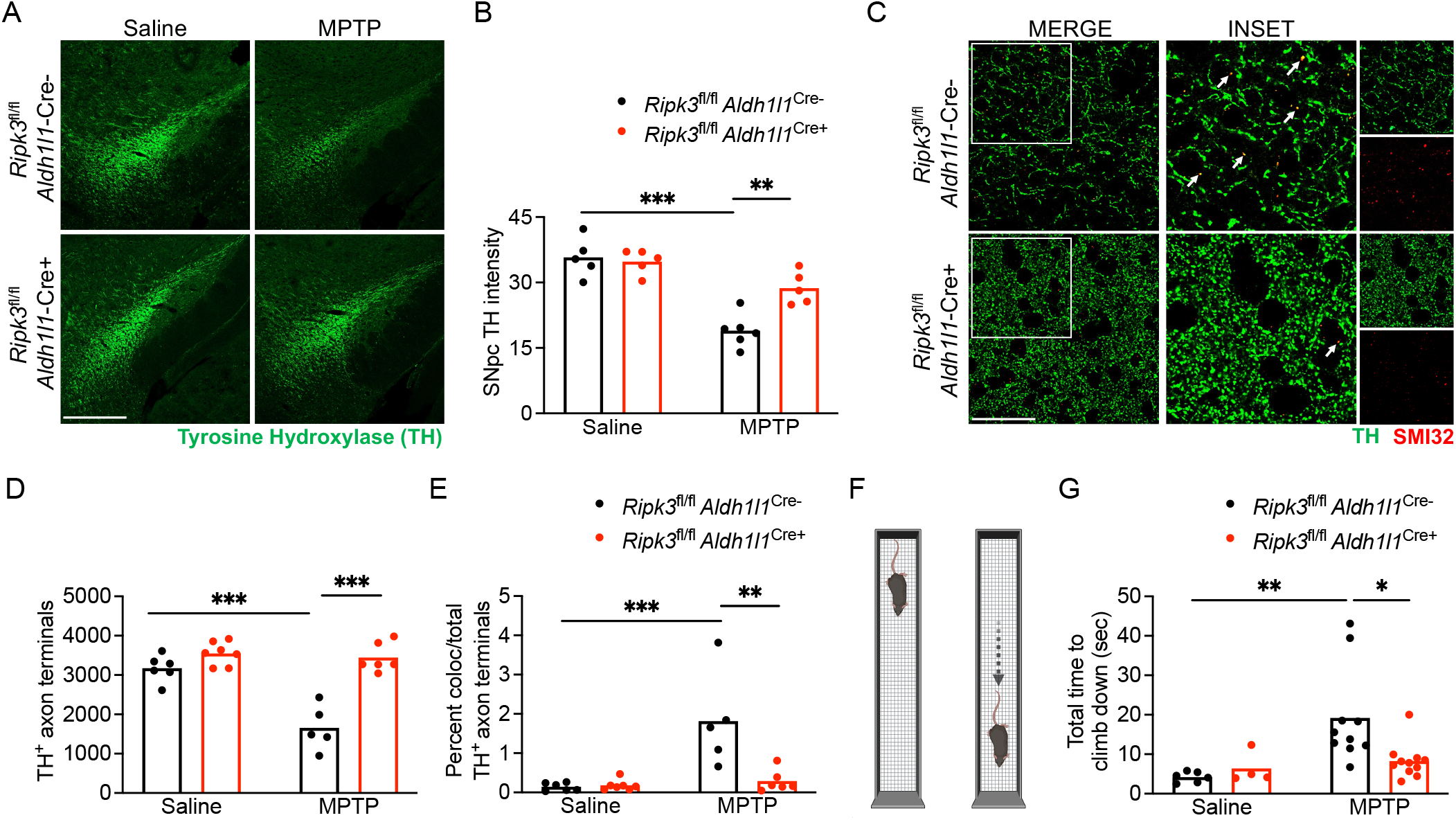
Astrocytic RIPK3 signaling promotes pathogenesis in the MPTP model of Parkinson’s disease. **(A-B)** IHC analysis of tyrosine hydroxylase (TH) staining in the substantia nigra pars compacta (SNpc) in indicated genotypes 7 days following either saline or MPTP treatment (scale bar = 200 μm). **(C)** IHC analysis of TH^+^ axons with colabeling with the damaged axon marker SMI-32 in the striatum in indicated genotypes 7 days following either saline or MPTP treatment (scale bar = 20 μm). **(F)** Schematic diagram for the vertical grid test. **(G)** Behavioral performance in the vertical grid test 6 days after injection with MPTP or saline. *p<0.05, **p < 0.01, ***p < 0.001. See also Figure S1.

### RIPK3 drives inflammatory transcriptional activation but not proliferation in midbrain astrocytes

Given these findings, we next questioned how RIPK3 signaling influences the phenotype of astrocytes in the setting of MPTP administration. Immunohistochemical (IHC) staining of SNpc sections revealed increased GFAP staining in MPTP-treated control animals, consistent with astrocyte activation, and this effect was blocked in *Ripk3*^fl/fl^ *Aldh1l1*^Cre+^ mice (Figure 2A-B). To test whether enhanced GFAP staining indicated proliferative astrogliosis, we performed flow cytometric analysis of astrocytes in the midbrain of MPTP-treated animals, which revealed no differences in GLAST^+^ astrocytes between genotypes (Figure 2C-D). These data suggested that enhanced GFAP staining was not due to increased numbers of astrocytes following MPTP administration, but rather a change in the astrocyte activation status. To test this idea, we performed qRT-PCR analysis of a panel of transcripts that we and others have shown to be associated with neurotoxic astrocyte activation in models of Parkinson’s disease^14,36,37^. We observed upregulation of 10 out of 14 transcripts in our analysis panel in midbrain homogenates derived from MPTP-treated littermate controls, while this activation signature was essentially abolished in *Ripk3*^fl/fl^ *Aldh1l1*^Cre+^ mice (Figure 2E). In contrast, MPTP-treated *Mlkl*^-/-^ mice showed equivalent levels of inflammatory transcript expression in the midbrain (Supplemental Figure 2). These data suggest that astrocytic RIPK3 signaling promotes an inflammatory transcriptional state in the midbrain following MPTP treatment, independently of MLKL and necroptosis.

**Figure 2.**
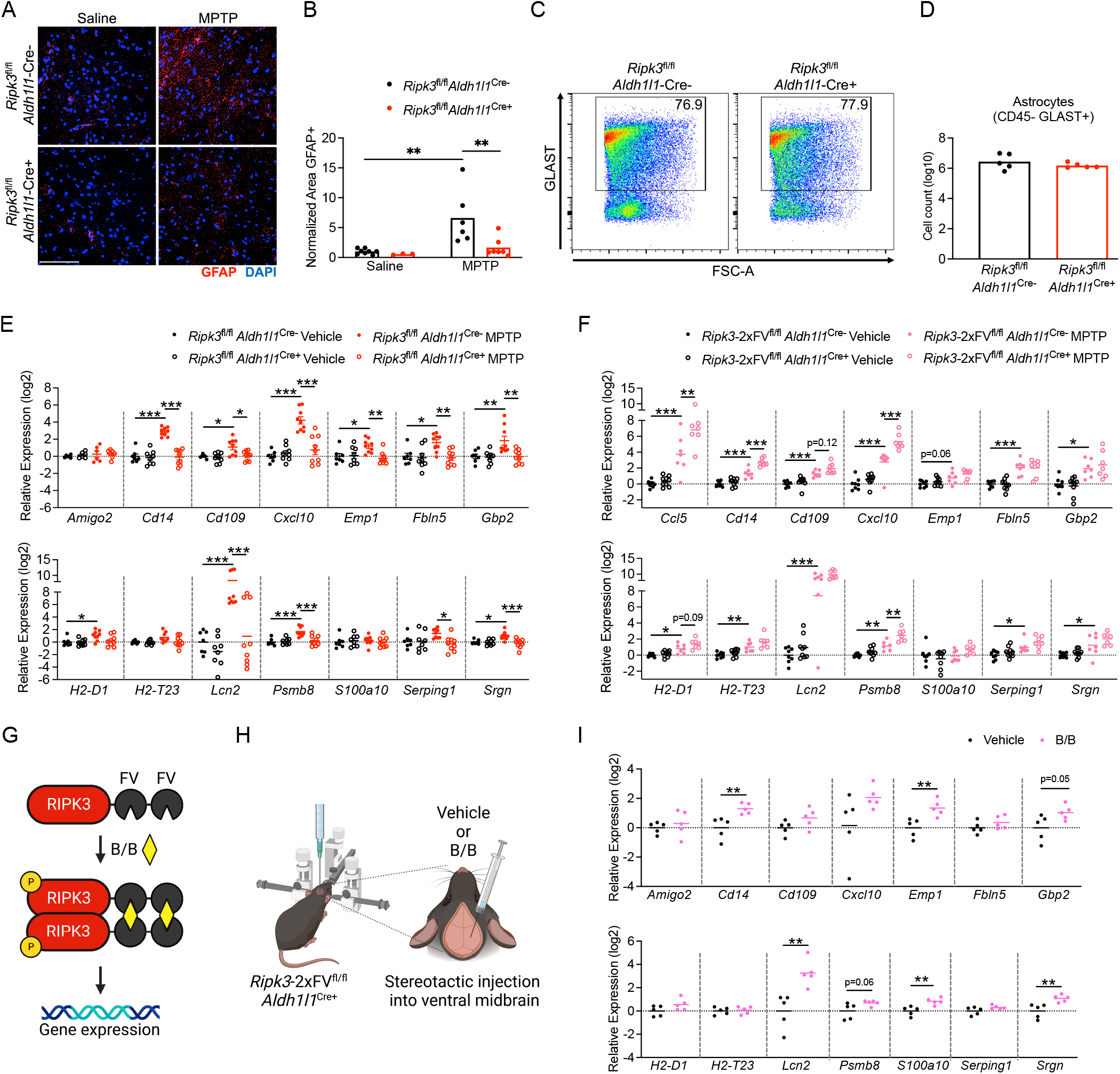
RIPK3 drives inflammatory transcriptional activation but not proliferation in midbrain astrocytes. **(A-B)** IHC analysis of GFAP staining in the substantia nigra pars compacta (SNpc) in indicated genotypes 3 days post-MPTP treatment (scale bar = 200 μm). **(C-D)** Flow cytometric analysis of GLAST+ astrocytes in midbrain homogenates derived from indicated genotypes 3 days post-MPTP treatment. **(E-F)** qRT-PCR analysis of indicated genes in midbrain homogenates derived from astrocyte-specific *Ripk3* knockouts **(E)** or astrocyte-specific *Ripk3* overexpressing **(F)** mice 3 days post-MPTP treatment. **(G-H)** Schematic of inducible RIPK3 activation system **(G)** and stereotactic delivery of dimerization drug into the ventral midbrain **(H)**. **(I)** qRT-PCR analysis of indicated genes in midbrain homogenates derived from *Ripk3*-2xFV^fl/fl^ *Aldh1l1*-Cre+ mice 24 hours following administration of B/B homodimerizer or vehicle control. *p<0.05, **p < 0.01, ***p < 0.001. See also Figure S2.

We next more carefully assessed this idea by using a mouse line expressing RIPK3 fused to two FKBP^F36V^ domains that facilitate enforced oligomerization following treatment with a dimerization drug. This protein is expressed in a cell type-specific manner under the control of a lox-STOP-lox element in the *Rosa26* locus, while the endogenous *Ripk3* locus is left intact. Thus, this mouse line can be used as both a cell type-specific overexpression system while also facilitating forced chemogenetic activation of RIPK3 in cell types of interest *in vivo*^12,13,38^. We first questioned whether simple overexpression of RIPK3 in astrocytes would enhance the inflammatory transcriptional signature that occurs following MPTP administration. We observed that 4 neurotoxic astrocyte-associated transcripts exhibited augmented upregulation following MPTP administration in *Ripk3*-2xFV^fl/fl^ *Aldh1l1*^Cre+^ mice, including *Ccl5, Cd14, Cxcl10,* and *Psmb8*, while 2 others exhibited trends towards increased expression that did not reach statistical significance (*Cd109, H2-D1*) (Figure 2F). To assess whether activation of astrocytic RIPK3 was sufficient to induce an inflammatory gene signature, we enforced RIPK3 activation in astrocytes via stereotactic delivery of B/B homodimerizer to the ventral midbrain of *Ripk3*-2xFV^fl/fl^ *Aldh1l1*^Cre+^ mice. B/B homodimerizer binds in a multivalent fashion to the FKBP^F36V^ domains of RIPK3-2xFV proteins, driving their oligomerization, which is sufficient to induce RIPK3 kinase activity in the absence of any other stimulus^39,40^ (Figure 2G-H). Enforced activation of RIPK3 in midbrain astrocytes *in vivo* resulted in induced expression of several neurotoxic astrocyte-associated transcripts, including *Cd14, Emp1, Gbp2, Lcn2, S100a10,* and *Srgn* (Figure 2I). Together, these data show that activation of RIPK3 in midbrain astrocytes drives their activation and the establishment of an inflammatory transcriptional signature.

### Astrocytic RIPK3 signaling has minimal impact on microgliosis in the MPTP model

We next questioned whether the reduced expression of inflammatory genes observed in mice lacking astrocytic RIPK3 was associated with cell non-autonomous effects on other cell types in the setting of MPTP treatment. We thus performed IHC staining for IBA1, a marker of myeloid cells that largely labels microglia in the setting of sterile neurodegeneration^41,42^. This analysis revealed no differences in the overall coverage of IBA1 staining in the midbrain in *Ripk3*^fl/fl^ *Aldh1l1*^Cre+^ mice compared to littermate controls, though IBA1^+^ cells did appear to exhibit a somewhat less ramified and more “activated” morphology following MPTP treatment in controls, but not conditional knockout, animals (Figure 3A-B). To assess changes to immune cells more carefully, we next performed flow cytometric analysis of leukocytes derived from the midbrain of MPTP-treated mice. This revealed essentially identical frequencies of CD45^int^ CD11b^+^ F4/80^+^ microglia between genotypes (Figure 3C-D), confirming a lack of difference in microglial proliferation. Microglia exhibited no differences in common activation markers, including MHC-II (data not shown), between genotypes, although microglia derived from MPTP-treated *Ripk3*^fl/fl^ *Aldh1l1*^Cre+^ mice exhibited diminished expression of the costimulatory molecule CD80 compared to controls (Figure 3E-F), consistent with a less inflammatory phenotype. We observed very low frequencies of CD45^hi^ infiltrating peripheral immune cells in the MPTP model (Figure 3C), the overall numbers of which did not differ by genotype (Figure 3G). These data suggest that astrocytic RIPK3 signaling following MPTP administration likely induces neuroinflammation primarily through cell-intrinsic mechanisms, with modest cell non-autonomous effects on microglia.

**Figure 3.**
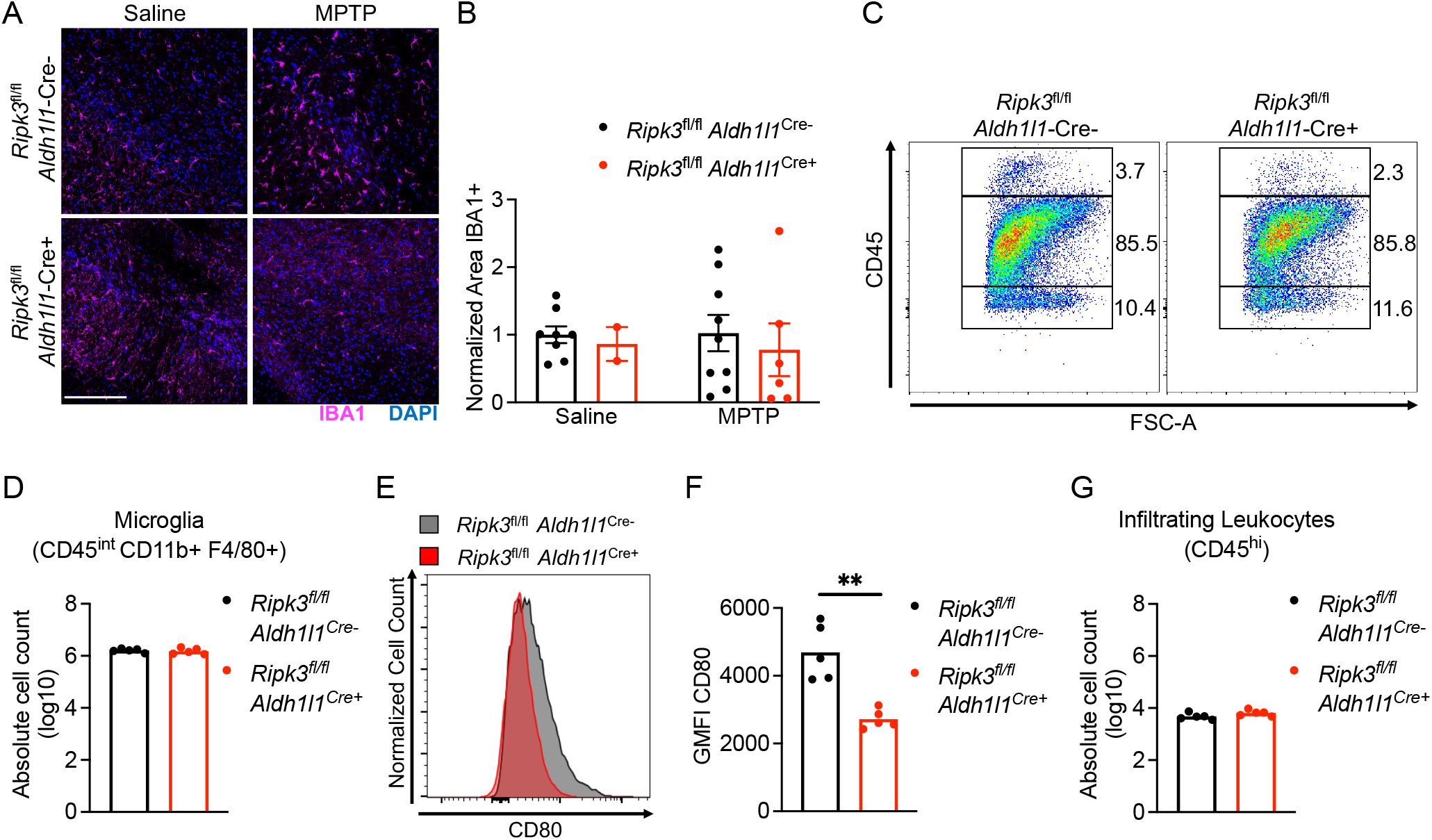
Astrocytic RIPK3 signaling has minimal impact on microgliosis in the MPTP model. **(A-B)** IHC analysis of IBA1 staining in the substantia nigra pars compacta (SNpc) in indicated genotypes 3 days post-MPTP treatment (scale bar = 200 μm). **(C)** Representative flow cytometric plot depicting leukocyte populations in midbrain homogenates derived from indicated genotypes 3 days pos-MPTP treatment. **(D)** Quantification of absolute numbers of microglia derived from flow cytometric analysis. **(E-F)** Representative histogram **(E)** and quantification of geometric mean fluorescence intensity (GMFI) **(F)** derived from analysis of CD80 expression on microglial populations in **(D)**. **(G)** Quantification of absolute numbers of CD45^hi^ leukocytes derived from flow cytometric analysis. **p < 0.01

### Astrocytic RIPK3 activation drives a transcriptomic state associated with inflammation and neurodegeneration in the midbrain

To characterize how astrocytic RIPK3 shapes the neuroinflammatory state of the brain more thoroughly in the MPTP model, we also performed bulk RNA sequencing (RNA-seq) of isolated midbrain tissues derived from *Ripk3*^fl/fl^ *Aldh1l1*^Cre+^ and littermate controls. Principle component analysis revealed distinct separation of MPTP-treated control animals along PC1, while MPTP-treated conditional knockouts largely clustered with vehicle-treated animals of both genotypes (Figure 4A). Further analysis revealed a robust transcriptional response to MPTP in midbrain tissues of littermate control animals, including 452 significantly upregulated genes and 145 significantly downregulated genes (Figure 4B) compared to vehicle-treated controls. This transcriptional response was blunted in *Ripk3*^fl/fl^ *Aldh1l1*^Cre+^ mice, which exhibited only 195 significantly upregulated genes and 120 significantly downregulated genes compared to genotype-matched vehicle-treated animals (Figure 4C), suggesting that astrocytic RIPK3 signaling drives a major portion of the tissue-wide transcriptional response to MPTP-induced neuronal cell death. In support of this idea, comparison of differentially expressed genes (DEGs) within MPTP-treated groups revealed 120 genes with significantly higher expression and 252 genes with significantly lower expression in conditional knockouts compared to littermate controls (Figure 4D).

**Figure 4.**
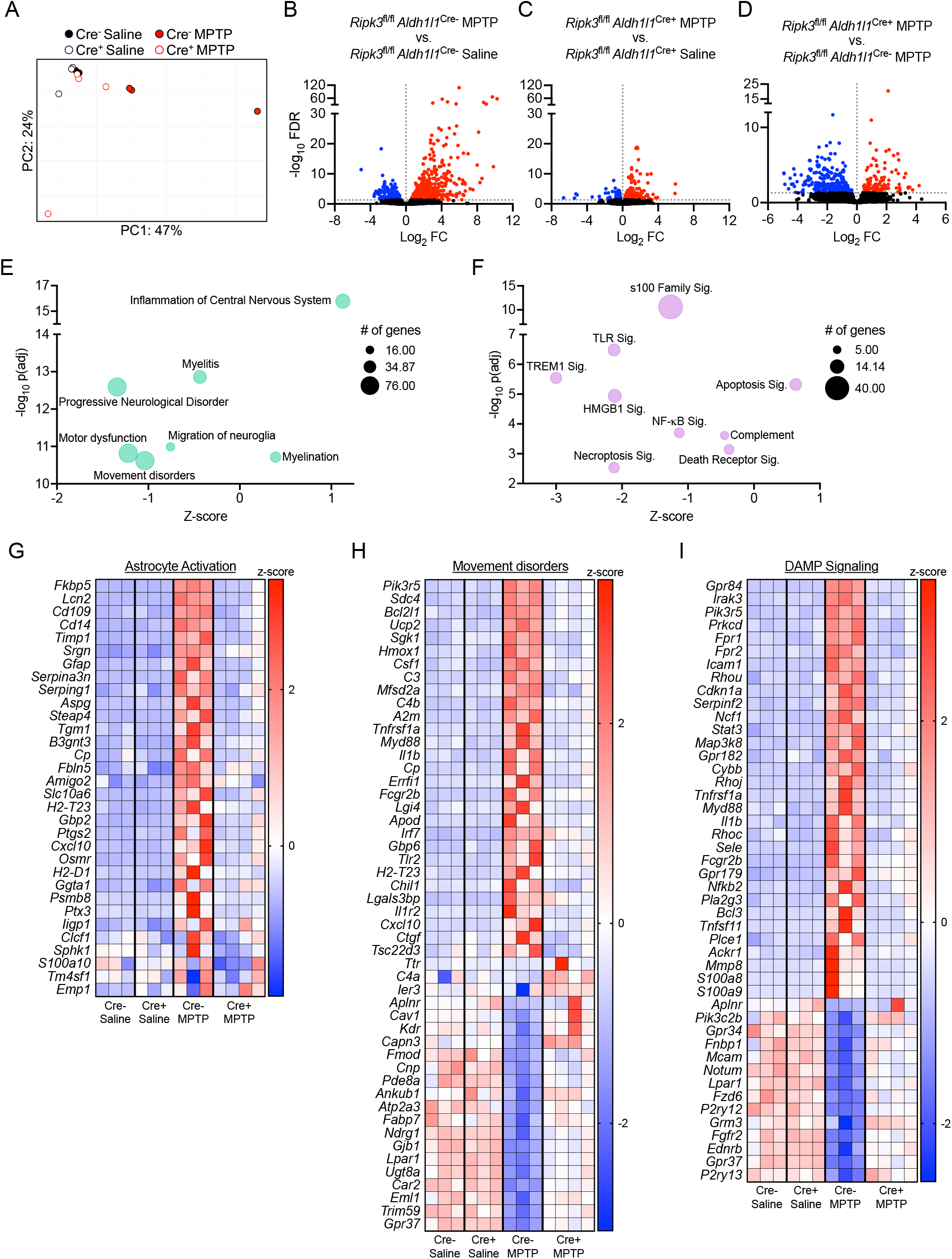
Astrocytic RIPK3 activation drives a transcriptomic state associated with inflammation and neurodegeneration in the midbrain. **(A-I)** Midbrains were harvested from mice of indicated genotypes 3 days post-treatment with MPTP or saline and subjected to bulk RNA-seq. (A) Principal component analysis demonstrating separation of treatment groups and genotypes in the RNA-seq dataset. (B-D) Volcano plots showing differentially expressed genes derived from indicated comparisons. Data points in red are genes exhibiting upregulated expression, while those in blue exhibit downregulated expression. Genes with an FDR <0.05 were considered significant. (E-F) Bubble plots showing selected significantly enriched disease and function terms (E) or canonical pathways (F) derived from Ingenuity Pathway Analysis comparing Cre- vs. Cre+ MPTP-treated groups. (G-I) Heatmaps showing significantly differentially expressed genes for selected pathways.

To better understand the functional relevance of these transcriptomic profiles, we performed Ingenuity Pathway Analysis (IPA) of genes differentially expressed between genotypes in MPTP-treated animals. This revealed significant enrichment of several disease and function terms with relevance to our study, including “Inflammation of the Central Nervous System,” “Progressive Neurological Disorder,” “Movement Disorders,” and others (Figure 4E). Comparisons of differentially regulated canonical pathways showed significant enrichment of pathways relating to programmed cell death and inflammation, as expected (Figure 4F). Notably, terms related to DAMP signaling were also highly enriched, including signaling by HMGB1 and S100 family proteins, both of which are factors released by dying and damaged cells that induce inflammation. Further analysis revealed significant upregulation of genes associated with astrocyte activation (Figure 4G), consistent with our previous qRT-PCR analysis. Comparisons of individual gene expression profiles for 2 selected IPA terms (Movement Disorders and DAMP signaling) revealed dozens of significant DEGs for both terms, characterized by a mix of both up-and down-regulated gene expression. Together, our RNA-seq analysis reveals a central role for astrocytic RIPK3 in promoting gene expression associated with neurodegeneration and neuroinflammation in the midbrain. Our findings also suggest a strong link between DAMP signaling and RIPK3-dependent neuroinflammation.

### Secreted factors from dying neurons drive RIPK3-dependent astrocyte activation

Given the strong representation of DAMP signaling in our transcriptomic analysis, we questioned whether factors released from dying neurons were important for driving RIPK3-mediated astrocyte activation. To test this, we treated differentiated SH-SY5Y neuroblastoma cells, a commonly used model of catecholaminergic neurons^43^, with the toxic MPTP metabolite MPP^+^ (5mM) for 24 hours, which resulted in around 50% cell death (Supplemental Figure 3A). We harvested the conditioned media (NCM) from these cells, which contained DAMPs and other factors released from dying SH-SY5Y cells, and added it to primary human midbrain astrocyte cultures at a ratio of 1:1 with normal astrocyte culture media (Figure 5A). NCM-treated astrocytes were also treated with the RIPK3 inhibitor GSK872 or DMSO vehicle. qRT-PCR analysis following 24 hours of stimulation under these conditions revealed robust induction of genes associated with inflammatory activation in midbrain astrocyte cultures treated with NCM derived from MPP^+^-treated SH-SY5Y cultures, hereafter referred to as MPP^+^ NCM (Figure 5B). However, pharmacologic inhibition of RIPK3 signaling in astrocytes largely prevented this effect.

**Figure 5.**
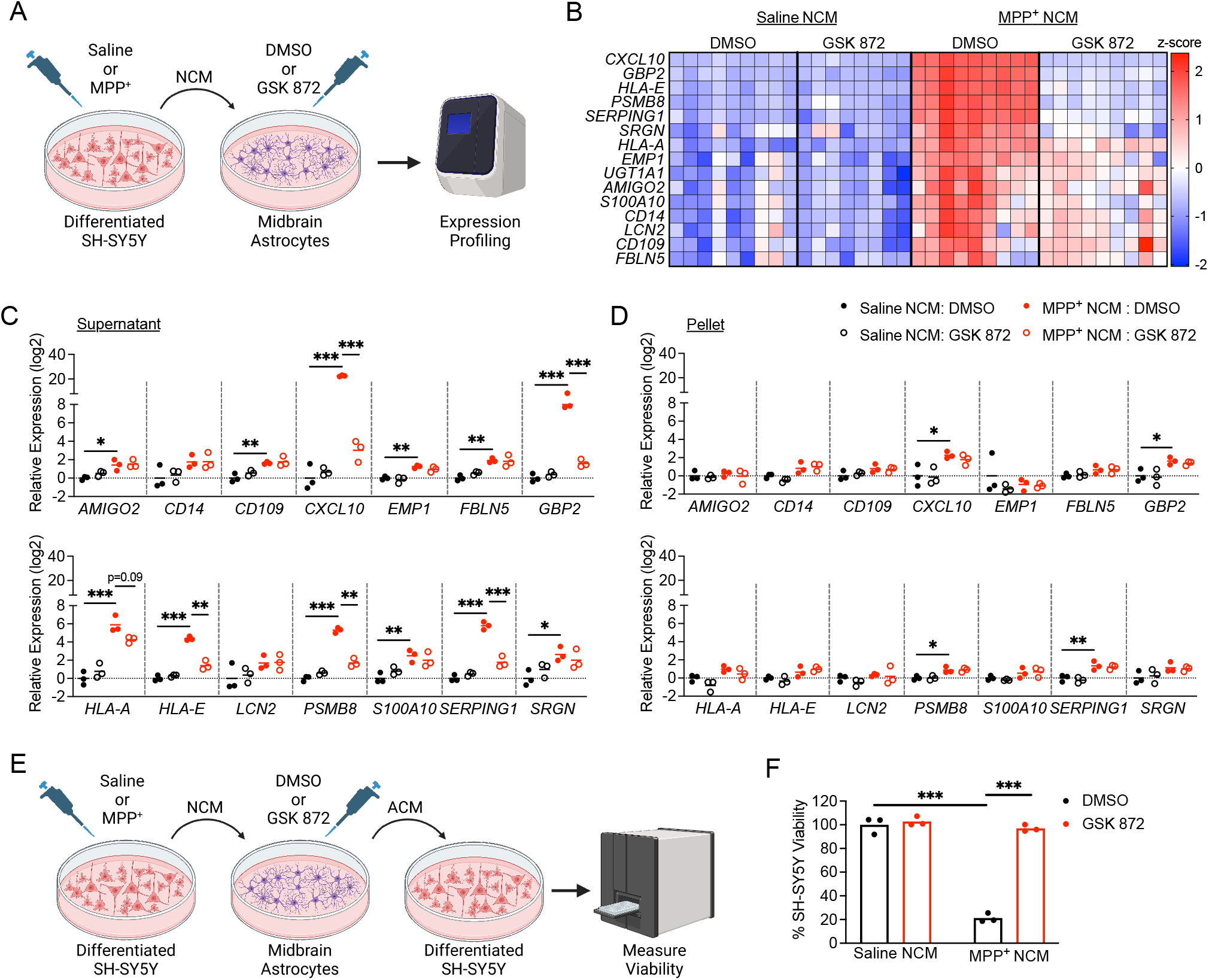
Secreted factors from dying neurons drive RIPK3-dependent astrocyte activation. **(A)** Schematic of experimental design for DAMP transfer experiments. Differentiated SH-SY5Y cells were treated with MPP^+^ or saline for 24h and media (NCM) was then transferred to cultures of primary human midbrain astrocytes. Astrocytes were treated with NCM in the presence of GSK 872 or control for 24h prior to qRT-PCR profiling. **(B)** Heatmap showing expression of astrocyte activation-associated genes in astrocyte cultures treated as in **(A)**. **(C-D)** qRT-PCR profiling of indicated genes in astrocytes treated for 24h with clarified NCM supernatants **(C)** or pelleted SH-SY5Y debris **(D). (E)** Schematic of experimental design for neurotoxicity assay. Astrocytes were treated with NCM as in **(A)** for 24h. Astrocytes were then washed and media replaced for another 24h. This new astrocyte conditioned medium (ACM) was then transferred to fresh SH-SY5Y cells for cell viability measurement. **(F)** Cell Titer Glo analysis of SH-SY5Y viability 24h following treatment with ACM derived from indicated conditions. *p<0.05, **p < 0.01, ***p < 0.001. See also Figures S3 and S4.

After these observations, we recognized that our NCM preparations may have contained debris and floating “corpses” from dead SH-SY5Y cells. To assess whether soluble factors or dead cell-associated material was the primary driver of RIPK3-dependent astrocyte activation in our experiments, we carefully fractionated NCM samples to pellet out cellular material from soluble factors in the media. Application of either clarified supernatant (Figure 5C) or resuspended pellet material (Figure 5D) from MPP^+^-treated SH-SY5Y cells to midbrain astrocyte cultures revealed that clarified supernatants stimulated expression of many inflammatory genes in astrocytes in a largely RIPK3-dependent manner. In contrast, pellet-derived material was only minimally stimulatory, and this stimulation was RIPK3-independent. We also confirmed that exposure to residual MPP^+^ in NCM was not the primary driver of astrocyte activation, as direct application of MPP^+^ to midbrain astrocyte cultures did not result in either cell death or upregulation of inflammatory gene expression (Supplemental Figure 3B-C).

We next wanted to confirm that inflammatory gene expression in our system corresponded to a functional readout of astrocyte activation. We thus assessed whether exposure to dying neuron-derived factors would confer neurotoxic activity to astrocytes. We first treated human midbrain astrocytes for 24 hours with MPP^+^ NCM with or without RIPK3 inhibitor (and respective controls), then washed the cells and replaced the astrocyte medium to remove residual MPP^+^. We then cultured astrocytes for an additional 24h and collected their conditioned media (ACM), which was then added to fresh cultures of SH-SY5Y cells at a 1:1 ratio with normal SH-SY5Y media (Figure 5E). We confirmed that astrocytes maintained transcriptional activation for at least 24 hours following this wash step, confirming that astrocytes remain activated after removal of MPP^+^ NCM in this paradigm (Supplemental Figure 4). ACM derived from MPP^+^ NCM-treated astrocytes induced around 80% cell death in fresh SH-SY5Y cultures after 24 hours, while this neurotoxic activity was completely abrogated when astrocytic RIPK3 signaling was inhibited (Figure 5F). Together, these data show that soluble factors released from dying neuron-like cells are sufficient to induce inflammatory transcription and neurotoxic activity in midbrain astrocytes and that this process requires, to a large degree, cell-intrinsic RIPK3 activity within astrocytes.

### DAMP signaling via RAGE drives inflammatory activation in midbrain astrocytes

We next sought to more precisely identify specific DAMP signals that stimulate midbrain astrocyte activation. Our transcriptomic analysis revealed that both HMGB1 and S100 family signaling were highly enriched in an astrocytic RIPK3-dependent manner in the midbrain following MPTP treatment. As both of these DAMPs stimulate a common receptor, RAGE, we assessed whether RAGE was required for astrocyte activation following exposure to MPP^+^ NCM. We thus treated human midbrain astrocyte cultures with MPP^+^ or control NCM, along with the RAGE inhibitor FPS-ZM1 for 24 hours and performed qRT-PCR profiling (Figure 6A). Blockade of RAGE in astrocytes substantially reduced MPP^+^ NCM-induced transcriptional activation, effectively preventing upregulation of 6 out of 11 astrocyte activation-associated transcripts (Figure 6B). Based on these findings, we confirmed that the RAGE ligand HMGB1 was, in fact, released by SH-SY5Y cells following induction of cell death by MPP^+^ (Figure 6C). We also observed significant accumulation of HMGB1 protein in midbrain homogenates of mice treated with MPTP (Figure 6D), confirming that induction of dopaminergic cell death results in the release of RAGE ligands *in vivo*. To assess whether RAGE ligands induced astrocyte activation in a RIPK3-dependent manner, we next treated primary midbrain astrocytes with recombinant DAMPs and profiled gene expression. Strikingly, we observed that stimulation of murine midbrain astrocytes with HMGB1 induced robust transcriptional activation that was blocked in the presence of GSK 872. As a complimentary approach, we also generated midbrain astrocyte cultures from *Ripk3^-/-^*mice (and heterozygous littermate controls) and stimulated with RAGE ligands. Treatment with either HMGB1 (Figure 6F) or S100β (Figure 6G) induced inflammatory transcript expression in control but not *Ripk3*^-/-^ cultures. Together, these data suggest that dying neurons release DAMPs that induce inflammatory astrocyte activation through activation of astrocytic RAGE, which in turn drives transcription via RIPK3 signaling.

**Figure 6.**
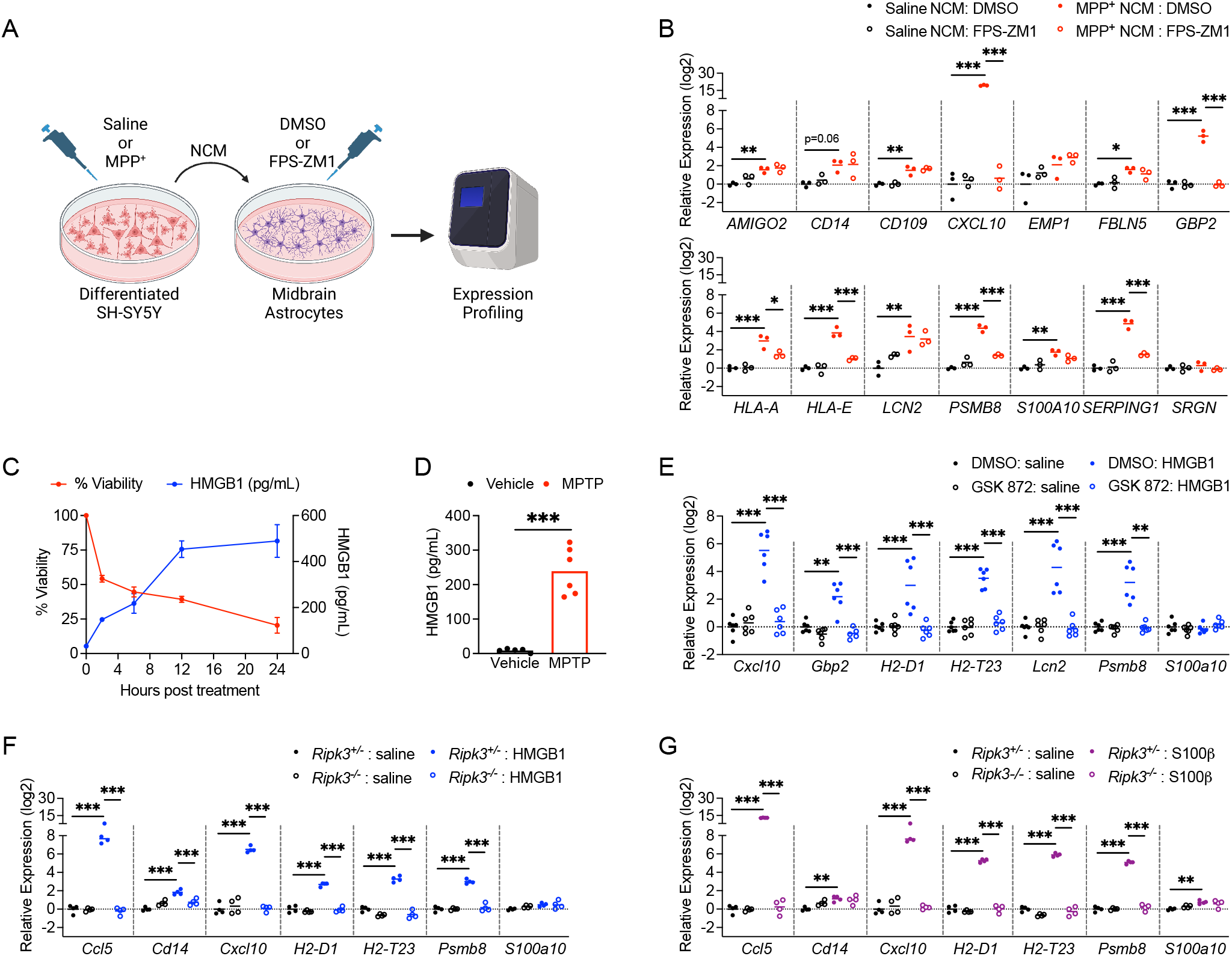
DAMP signaling via RAGE drives inflammatory activation in midbrain astrocytes. **(A)** Schematic of experimental design for DAMP transfer experiments. Differentiated SH-SY5Y cells were treated with MPP^+^ or saline for 24h and media (NCM) was then transferred to cultures of primary human midbrain astrocytes. Astrocytes were treated with NCM in the presence of FPS-ZM1 or control for 24h prior to qRT-PCR profiling. **(B)** qRT-PCR profiling of indicated genes in astrocytes treated for 24h with NCM derived from indicated conditions**. (C-D)** ELISA analysis of HMGB1 protein levels in supernatants of SH-SY5Y cells treated with MPP^+^ **(C)** or midbrain homogenates from WT mice 3 days post-MPTP treatment **(D)** n=4-8 replicates per time point in **(C). (E-G)** qRT-PCR analysis of indicated genes in WT murine midbrain astrocytes **(E)** or midbrain astrocytes derived from indicated genotypes **(F-G)** 24h following treatment with recombinant HMGB1 **(E-F)** or S100β **(G).** *p<0.05, **p < 0.01, ***p < 0.001.

### Activation of RIPK3 by DAMP signaling drives pathogenic functional changes in midbrain astrocytes

To confirm that the transcriptional effects of DAMP signaling impacted astrocyte function, we collected astrocyte conditioned media (ACM) from astrocytes treated for 24h with MPP^+^ NCM with or without RAGE inhibitor (and respective controls) and applied the ACM to fresh cultures of SH-SY5Y cells (Figure 7A). ACM derived from MPP^+^ NCM-treated astrocytes induced significant cell death in fresh SH-SY5Y cultures, while this neurotoxic activity was completely abrogated when astrocytic RAGE signaling was inhibited (Figure 7B). We also observed conferral of neurotoxic activity following direct stimulation of astrocytes with recombinant DAMPs (Figure 7C), including HMGB1 (Figure 7D) and S100β (Figure 7E). However, this neurotoxic activity was also abrogated when RIPK3 signaling was blocked, further supporting a role for a RAGE-RIPK3 axis in promoting neurotoxic astrocyte activation. This neurotoxic activity was not due to residual recombinant DAMPs in ACM, as direct application of either DAMP ligand to SH-SY5Y cells did not result in cell death (Supplemental Figure 5). As previous work has shown that neurotoxic astrocytes downregulated key homeostatic functions such as phagocytosis^14,36^, we also exposed midbrain astrocyte cultures to labeled debris generated from SH-SY5Y cells and measured phagocytic uptake of debris via flow cytometry (Figure 7F). Direct stimulation of astrocytes with HMGB1 resulted in a significant reduction in uptake of CSFE-labeled debris, while this suppression of phagocytic function was blocked in the presence of a RIPK3 inhibitor (Figure 7G-H). We also observed that MPP^+^ NCM similarly reduced astrocytic phagocytosis in a RIPK3-dependent fashion (Figure 7I). These data further support the notion that DAMPs emanating from dying neurons alter astrocytic function via activation of RIPK3 signaling.

**Figure 7.**
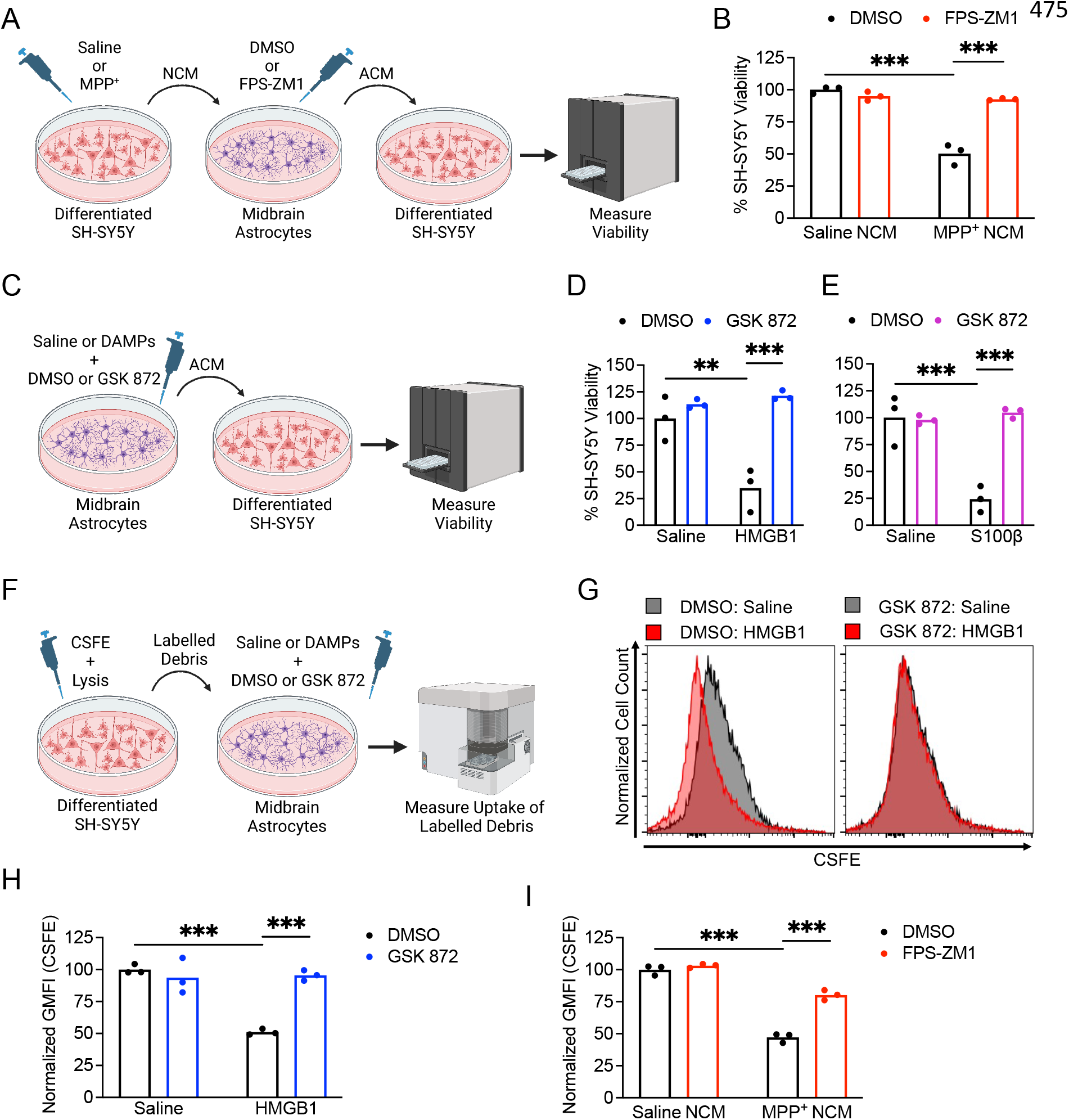
Activation of RIPK3 by DAMP signaling drives pathogenic functional changes in midbrain astrocytes. **(A)** Schematic of experimental design for neurotoxicity experiments. Differentiated SH-SY5Y cells were treated with MPP^+^ or saline for 24h and media (NCM) was then transferred to cultures of primary human midbrain astrocytes. Astrocytes were treated with NCM in the presence of FPS-ZM1 or control for 24h. Astrocytes were then washed and media replaced for another 24h. This new astrocyte conditioned medium (ACM) was then transferred to fresh SH-SY5Y cells for cell viability measurement. **(B)** Cell Titer Glo analysis of SH-SY5Y viability 24h following treatment with ACM derived from indicated conditions. **(C)** Schematic showing treatment of primary human midbrain astrocytes with recombinant DAMPs for 24h prior to transfer of ACM to SH-SY5Y cultures and measurement of cell viability. **(D)** Cell Titer Glo analysis of SH-SY5Y viability 24h following treatment with ACM derived from indicated conditions**. (F)** Schematic showing generation and transfer of CSFE-labeled neuronal debris to midbrain astrocytes treated with recombinant DAMPs with or without GSK 872. Astrocytes were cultured in the presence of labelled debris for 24h and then CSFE internalization was measured via flow cytometry. **(G-H)** Representative histograms **(G)** and quantification of GMFI **(H)** of CSFE signal in astrocytes treated as in **(F). (I)** GMFI of CSFE internalization in astrocytes treated as in **(F)** but with NCM rather than recombinant DAMPs and FPS-ZM1 rather than GSK 872. **p < 0.01, ***p < 0.001. See also Figure S5.

## Discussion

Our study defines a previously unknown role for neuronal DAMPs in promoting neurotoxic astrocyte activation. This effect was mediated by RIPK3-mediated transcriptional activation, an effect that occurred independently of the necroptotic executioner protein MLKL. Mechanistically, we found that astrocytic RAGE signaling was required for astrocyte activation downstream of DAMP exposure, and this RAGE/RIPK3 signaling axis promoted inflammatory transcription and neurotoxic functional activity. Intriguingly, these results suggest that neuronal death, itself, potentiates a feed-forward process of astrocyte activation and further neuronal cell death. These findings highlight an important mechanism of neuron-glia crosstalk in the pathogenesis of neurodegeneration.

DAMPs have previously been implicated as drivers of inflammation in a broad variety of disorders, including neurodegeneration, ischemic stroke, autoimmunity, cardiovascular disease, and others^44-55^. RAGE ligands, in particular, have been associated with neurodegenerative disease and have been the target of preclinical therapeutic development. For example, S100β levels in serum and cerebrospinal fluid (CSF) has been shown to correlate with disease severity in Parkinson’s disease^27,56^. Mice deficient in S100β are also resistant to MPTP-driven neurodegeneration^27^, consistent with a role for this molecule in perpetuating neuronal cell death. Similarly, antibody-mediated neutralization of HMGB1 has been shown to attenuate glial cell activation and prevent neuron loss in models of both Alzheimer’s disease and Parkinson’s disease^26,57^. Despite these findings, other groups have also described neuroprotective functions for RAGE ligands^58^, including stimulation of neurotrophic growth factor expression in amyotrophic lateral sclerosis^59^, suppression of amyloidosis^60^, and direct anti-apoptotic effects in neurons^61,62^. These complex effects appear to be highly context-dependent, differing by cell type, disease state, and even DAMP concentration^61,63,64^. Our data support a pathogenic role for RAGE signaling in the promotion of neurotoxic astrocyte activation.

Astrocytes express RAGE and other DAMP sensors, although cell type-specific functions for DAMP signaling in astrocytes have not been thoroughly studied^65^. Existing studies suggest that astrocytic RAGE signaling is pathogenic, on balance^66-68^. In Huntington’s disease, RAGE-positive astrocytes have been shown to have high levels of nuclear NF-κB^67^, consistent with a role for this pathway in promoting inflammatory astrocyte activation. Diminished levels of HMGB1 following berberine treatment was also correlated with diminished astrocyte activation in a model of sepsis^69^. Astrocytes are also major sources of RAGE ligands, particularly S100β, and much work to date has focused on autocrine RAGE signaling in astrocytes as a result^70-72^. We took advantage of the MPTP model, which induces death selectively in neurons but not astrocytes^73^, as well as serial culture systems to more directly assess the impact of paracrine RAGE signaling on astrocyte activation and function. Our study suggests that DAMPs released from dying neurons potently induce inflammatory astrocyte activation via RAGE, driving neurotoxic activation and perpetuating further neuronal cell death. These findings identify RAGE as a promising target for modulating astrocytic responses to neuronal cell death during neurodegenerative disease.

RIPK3 signaling has previously been shown to drive pathogenic neuroinflammation and neuronal cell death in several models of neurological disorders^14,15,74-77^. While many studies have reported neuronal necroptosis as a driver of neurodegeneration, we and others have described necroptosis-independent functions for this kinase in the coordination of neuroinflammation^12-18^. To date, RIPK3 signaling in astrocytes has received relatively little attention. Our findings here suggest that DAMP signaling activates astrocytic RIPK3 via RAGE signaling, which drives an inflammatory transcriptional program, even in the absence of MLKL. These data suggest that astrocytic RAGE signaling does not induce inflammation via necroptosis, consistent with our prior work showing necroptosis-independent RIPK3 signaling in astrocytes exposed to fibrillar α-synuclein^14^.

Future work will be needed to define the signaling events that mediate RAGE-dependent RIPK3 activation. A recent study demonstrated co-immunoprecipitation of RIPK3 with RAGE in an endothelial cell line following stimulation with TNF-α78, but the nature of this interaction and whether it happens under natural conditions *in vivo* remains to be established. While some studies have observed RIPK3 activation downstream of HMGB1^79,80^, these effects may have been mediated by non-RAGE HMGB1 receptors such as TLR4, which is known to stimulate RIPK3 via its adaptor molecule TRIF^81,82^. Both RAGE and RIPK3 signaling appear to converge on the potent activation of NF-κB^38,83-92^, which may provide clues concerning their potential molecular interactions. In any event, delineating the molecular events that promote pathogenic astrocyte activation downstream of DAMP signaling will likely be required to effectively target this pathway for future therapeutic development.

## Acknowledgements

The authors thank Drs. Noriko Goldsmith and Jessica Shivas for assistance with confocal imaging in the Human Genetics Institute of New Jersey Imaging Core Facility. We also thank Eric Chiles and Dr. Xiaoyang Su for assistance with LC/MS. LC/MS data were generated by the Rutgers Cancer Institute of New Jersey Metabolomics Shared Resource, supported, in part, with funding from NCI-CCSG P30CA072720-5923. Some figure elements were created with Biorender.com.

This work was supported by a research grant from the American Parkinson’s Disease Association, NIH R01 NS120895, and startup funds from Rutgers University (all to BPD), as well as R00 MH112855 (to RH). NPC was supported by F31 NS124242. WRE was supported by T32 AA028254. DA was supported by fellowships from the Louis Stokes Alliance for Minority Participation (LSAMP) program.

## Author Contributions

Conceptualization: NPC, BPD; Investigation: NPC, ED, WRE, MN, MM, DA, ML, TC, CA, BPD; Analysis: NPC, ED, MM, ML, TC, BPD; Resources: AWK, RH, BPD; Writing –Original Draft: NPC, BPD; Writing – Review and Editing: NPC, ED, TC, CA, BPD; Supervision: CA, AWK, RH, BPD; Funding Acquisition: RH, BPD.

## Competing Interests

The authors declare no competing interests.

## Methods

### Mouse lines

Mice were bred and housed under specific-pathogen free conditions in Nelson Biological Laboratories at Rutgers University. *Ripk3*^-/-^ and *Ripk3*^fl/fl^ mouse lines were generously provided by Genentech, Inc. *Mlkl*^-/-93^ and *Ripk3*-2xFV^fl/fl12^ lines were provided by Andrew Oberst (University of Washington). *Aldh1l1*-Cre/ERT2 mice were obtained from Jackson Laboratories (Line 031008) and all animals expressing this transgene were treated for five days with 60 mg/kg tamoxifen (Sigma-Aldrich, T5648) in sunflower oil (Sigma-Aldrich, S5007) (i.p.) at least one week prior to further experimentation. All genotyping was performed in house using ear punch tissue lysed overnight in DirectPCR Lysis Reagent (Viagen, 102-T) and Proteinase K (Sigma, #3115828001). Sequences for genotyping primers are listed in the Supplementary Table S1. PCR bands were visualized on 2% agarose (VWR, 97062) in TBE (VWR, E442) and stained in Diamond Nucleic Acid Stain (Promega, H1181). All experiments were performed in 8-12 week old animals, following protocols approved by the Rutgers University Institutional Animal Care and Use Committee (IACUC). All MPTP experiments were performed in male animals, as female animals experience high rates of toxicity and mortality in this model^29^. Other experiments, including B/B homodimerizer administration and primary cell culture, used balanced groups of both male and female animals.

### MPTP model

1-methyl-4-phenyl-1,2,3,6-tetrahydropyridine (MPTP) was administered at 20 mg/kg (i.p.) once per day for five days ^94^. Animals were harvested three days following the final MPTP injection for gene expression and flow cytometry experiments. Animals were harvested seven days after the last injection for immunofluorescence (IF) and vertical grid maze studies.

### Tissue collection

Mice were perfused transcardially with ice cold phosphate-buffered saline (PBS) followed by 4% paraformaldehyde (PFA) for IF experiments. Perfused brains were stored in 4% PFA overnight followed by 48 hours in 30% sucrose in PBS. For transcriptional and ELISA studies, mice were perfused with PBS and midbrain and/or striatal tissues were collected and homogenized for downstream analyses.

### Cell culture and treatment

Primary human midbrain astrocytes (ScienCell Research Laboratories) were cultured in astrocyte media (ScienCell, 1801) supplemented with 2% heat-inactivated fetal bovine serum (FBS) (ScienCell, 0010), astrocyte growth supplement (ScienCell, 1852), and penicillin/streptomycin (ScienCell, 0503). Cells from at least two distinct donors were used for all experiments. Human neuroblastoma SH-SY5Y cells (ATCC, CRL-2266) were cultured in DMEM medium (VWR, 0101–0500) supplemented with 10% FBS (Gemini Biosciences, 100–106), nonessential amino acids (Hyclone, SH30138.01), HEPES (Hyclone, 30237.01), penicillin/streptomycin (Gemini Biosciences, 400–110), and amphotericin B antifungal (Gemini Biosciences, 100–104). Differentiation and experimentation occurred in stocks having undergone less than 15 passages. SH-SY5Y neuroblastoma cells were differentiated into mature neuron-like cells by treating with retinoic acid (4 μg/mL; Sigma-Aldrich, R2625) and BDNF (25 ng/mL; Sigma-Aldrich, B3795) in low serum (2%) SH-SY5Y media. Differentiated SH-SY5Y cultures were used for experiments five to seven days post-differentiation. MPP^+^ iodide (Sigma-Aldrich, D048) was formulated in water to a stock concentration of 500 mM. Recombinant HMGB1 (R&D Systems, 1690-HMB-050) and S100B (Human: R&D Systems, 1820-SB; Mouse: Novus Biologicals, NBP2-53070) were formulated according to manufacturer recommendations. For cell culture experiments, all recombinant DAMPs were used at a final concentration of 100 ng/mL for 24 h before collection of preconditioned media and cell lysates. GSK 872 was purchased from Millipore Sigma (530389). FPS-ZM1 was purchased from Sigma-Aldrich (55030). All inhibitors were solubilized in DMSO and used at a final concentration of 1 μM.

### Primary mouse astrocyte isolation and culture

Primary mouse midbrain astrocytes were cultured from dissected midbrain tissues derived from mouse pups on postnatal day three (P3). Tissue was dissociated using Miltenyi Neural Dissociation Kit (T) following manufacturer’s instructions (Miltenyi, 130-093-231). Midbrain astrocytes were cultured on fibronectin-coated flasks and non-astrocytic cells were removed via differential adhesion, as previously described^95^. Astrocytes were expanded in AM-a medium (ScienCell, 1831) supplemented with 10% FBS, Astrocyte Growth Supplement-animal (ScienCell, 1882) and Penicillin/Streptomycin Solution (ScienCell, 0503).

### Cell viability test

Cell viability was assessed with the CellTiter-Glo Luminescent Cell Viability Assay kit (Promega, G7573), according to the manufacturer’s instructions. Luminescence signal was measured with a SpectraMax iD3 plate reader (Molecular Devices).

### Phagocytosis assay

Differentiated SH-SY5Y neuronal cells were labeled with BioTracker CSFE Cell Proliferation Kit (Millipore Sigma, SCT110) according to the manufacturer’s protocol. Cell death was induced by exposure to TNF-α at 100 ng/mL and cycloheximide (Sigmal-Aldrich, 66-81-9) at 100 μg/mL for 24 h. Labelled cell debris was collected by centrifugation. Unlabeled neuronal debris was used as a staining control. To detect phagocytosis, CSFE-labeled neuronal debris was added to primary midbrain astrocyte cultures at a ratio of 1:100 for 24 h. Excess neuronal debris was washed away with PBS. Astrocytes were then harvested with cold 5 mM EDTA in PBS followed by scraping of adherent cells. Astrocytes were stained with Zombie NIR at 1:1000 in 1XPBS according to the manufacturer’s protocol, followed by fixation in 1% PFA. Phagocytosed CSFE signal was detected using a Northern Lights flow cytometer (Cytek). Analysis was performed by FlowJo software (FlowJo LLC).

### Immunofluorescence

Brains were cryosectioned at 12 μm per slice and mounted on a charged slide. Following thawing in a humidified chamber, tissues were incubated in blocking solution consisting of 5% goat serum (Gibco, 16210) and 0.2% Triton X-100 for one hour at room temperature. Sections were then incubated with primary antibody diluted in blocking solution overnight at 4°C in a humidified chamber. Slides were then washed three times with PBS for 15 minutes followed by incubation in secondary antibody diluted in blocking solution for one hour at room temperature. Slides were washed three times to remove secondary antibody and were then stained with 4’,6-diamindino-2-phenylindole (DAPI; Biotium, 40011) diluted in PBS for 20 minutes at room temperature, followed by another wash. Sections were cover-slipped with Prolong Diamond Antifade Mountant medium (Invitrogen, P36930). Slides were allowed to dry and images were acquired using Airyscan fluorescent confocal microscope (Carl Zeiss, LSM 800).

### B/B homodimerizer and stereotactic injection

B/B homodimerizer was purchased from Takara USA lnc. (AP20187) and was formulated according to manufacturer’s recommendations. Buprenorphine extended-release (3.25mg/kg) was administered subcutaneously immediately prior to surgery. Mice were anaesthetized with isoflurane (4% induction, 1% maintenance) and positioned on a heating pad while the head was fixed for stereotactic injection. Each animal received 500 nL of freshly formulated B/B homodimerizer or vehicle delivered by a glass pipette using a Programmable Nanoject III Nanoliter Injector (Drummond) unilaterally into the right ventral lateral midbrain (relative to bregma: coordinates A/P: -3.00mm, M/L: -1.20mm, D/V: -4.50mm). The scalp was sutured, and animals were allowed to recover for 24 h before transcriptional analyses.

### Quantitative real-time PCR

Total RNA from homogenized midbrain tissues was extracted using Zymo Direct-zol RNA Miniprep kit, following manufacturer’s instructions (Zymo, R2051). Total RNA from cultured cells was isolated using Qiagen RNeasy Mini Kit according to the manufacture’s protocol (Qiagen, 74106). RNA yield and quality of the samples were assessed using a NanoDrop spectrophotometer. cDNA was then synthesized with qScript cDNA Synthesis Kit (Quantabio, 95047), followed by qRT-PCR with SYBR Green Master Mix (Bio-Rad, 1725275). Cycle threshold (Ct) values were obtained using QuantStudio 5 instrument (Applied Biosystems). Delta Ct was calculated as normalized to Ct values of the housekeeping gene 18S (CtTarget − Ct18S = ΔCt). Z-scores were calculated to graph heatmaps. Primer sequences in our study are listed in Supplementary Table S2.

### Flow Cytometry

After perfusing with ice-cold PBS, mouse midbrains were dissected and minced with a blade. Tissues were then further homogenized via 30 minute incubation in pre-warmed digestion buffer consisting of 2% FBS, 1% glutamine, 1% non-essential amino acids, 1% penicillin/streptomycin/amphotericin, and 1.5% HEPES, with 0.7U/mL collagenase VIII and 50U/mL DNase I on an orbital shaker. Triturated tissue homogenate was then passed through a 70 μm cell strainer and centrifuged at 350*xg* at 4°C for 10 minutes to obtain a single-cell suspension. Cell gradient separation was then achieved by resuspending the pellet in 20% bovine-serum albumin (BSA) in DMEM followed by 20 minute centrifugation at 4°C. After removing the myelin layer, the cell gradient was disrupted by inverting in additional FACS buffer that consisted of 1mM EDTA in PBS with 1% BSA. Resuspended cells were then incubated in antibodies for 30 min at 4°C in the dark. After washing with cold FACS buffer, cold 1% paraformaldehyde was then used to fix the cells. Data collection and analysis were performed using a Cytek Northern Lights Cytometer and FlowJo software. Data were normalized using standard counting beads (ThermoFisher, #C36950).

### HMGB1 enzyme-linked immunosorbent assay (ELISA)

HMBG1 ELISA (Novus Biologicals, NBP2-62766) was performed following the manufacturer’s protocol.

### Liquid chromatography-mass spectrometry (LC-MS)

A single dosage of MPTP (40 mg/kg) was administered for LC-MS analysis of MPP^+^ *in vivo*. Mice were transcardially perfused with ice-cold PBS 90 min after MPTP injection. Whole brain tissues were then isolated and homogenized in CryoMill tubes containing cold 40:40:20 methanol:acetonitrile:water solution with 0.5% Formic Acid. Following a 10 min incubation on ice, tissue homogenates were then centrifuged in the cold room for 10 min for 16,000 *xg*. Supernatants were then transferred to a new collection tube. The final sample was then treated with 15% NH4HCO3. LC/MS was performed at the Metabolomics Shared Resource Core Facility at the Rutgers Cancer Institute of New Jersey (New Brunswick, NJ).

### Behavioral assessment

The vertical grid motor assessment task was adapted from previous work^34^. Briefly, mice were acclimated to the vertical grid apparatus 3 times a day for 2 consecutive days. On each day, each mouse was placed on the inside of the apparatus 3 cm from the top, facing upward, and was allowed to turn around and climb down. The trial was repeated whenever the mouse failed to climb down and/or turn around within 60 seconds. The same trials were repeated on the day following acclimation and video recorded for analysis.

### Bulk RNA sequencing

Total RNA from midbrain tissues was extracted and assessed as described above. RNA samples were sent to Azenta (Piscataway, NJ) for library preparation and Next Generation Sequencing. RNA yield and sample quality were assessed with Qubit (Invitrogen) and TapeStation (Agilent). The Illumina HiSeq platform and 2 x 150-bp paired-end reads were used for the RNA sequencing. Initial analysis was processed by Azenta. The quality of raw RNA-seq data (FASTQ) files were evaluated using FASTQC. Sequence reads were trimmed to remove possible adapter sequences and nucleotides with poor quality using Trimmomatic v.0.36. Trimmed reads were then mapped to the mouse reference genome (GRCm38) available on ENSEMBL using the STAR aligner v.2.5.2b. Unique gene hit counts were calculated by using featureCounts from the Subread package v.1.5.2. The gene hit counts table was used for downstream differential expression analysis via DESeq2. Further statistical analysis was performed using R.

### Image analysis

To quantify TH^+^ and SMI32^+^ puncta and co-localization, images were processed by Imaris software (Oxford Instruments, Bitplane 9.5). Object based co-localization was used with the “Coloc” feature. For TH^+^ and SMI32^+^ particles, the spot detection function was used to define particles by first creating ‘vesicles’ in each channel. Input intensity for threshold was chosen to best represent the signal for both channels. Colocalized particles were defined with the “classification” feature, where the distance between TH^+^ and SMI32^+^ particles within 1 μm or less is considered co-localization. The percentage area and mean intensity of GFAP^+^ and IBA1^+^ signal were assessed using Fiji (ImageJ) software.

### Statistical analysis

Statistical analysis was completed using GraphPad Prism 9 (GraphPad). Normally distributed data were analyzed using appropriate parametric tests: student’s t test (2-tailed) or two-way analysis of variance (ANOVA) with Tukey’s post hoc test used to determine significant differences between groups. A p value less than 0.05 was considered statistically significant. All data points represent biological replicates unless otherwise noted.

## Notes

### Competing Interest Statement

The authors have declared no competing interest.

## References

1. Gilhus, N.E., and Deuschl, G. (2019). Neuroinflammation - a common thread in neurological disorders. Nat Rev Neurol 15, 429–430. 10.1038/s41582-019-0227-8.

2. Boyd, R.J., Avramopoulos, D., Jantzie, L.L., and McCallion, A.S. (2022). Neuroinflammation represents a common theme amongst genetic and environmental risk factors for Alzheimer and Parkinson diseases. J Neuroinflammation 19, 223. 10.1186/s12974-022-02584-x.

3. Giovannoni, F., and Quintana, F.J. (2020). The Role of Astrocytes in CNS Inflammation. Trends Immunol 41, 805–819. 10.1016/j.it.2020.07.007.

4. Endo, F., Kasai, A., Soto, J.S., Yu, X., Qu, Z., Hashimoto, H., Gradinaru, V., Kawaguchi, R., and Khakh, B.S. (2022). Molecular basis of astrocyte diversity and morphology across the CNS in health and disease. Science 378, eadc9020. 10.1126/science.adc9020.

5. Patani, R., Hardingham, G.E., and Liddelow, S.A. (2023). Functional roles of reactive astrocytes in neuroinflammation and neurodegeneration. Nat Rev Neurol 19, 395–409. 10.1038/s41582-023-00822-1.

6. Escartin, C., Galea, E., Lakatos, A., O’Callaghan, J.P., Petzold, G.C., Serrano-Pozo, A., Steinhauser, C., Volterra, A., Carmignoto, G., Agarwal, A., et al. (2021). Reactive astrocyte nomenclature, definitions, and future directions. Nat Neurosci 24, 312–325. 10.1038/s41593-020-00783-4.

7. Brandebura, A.N., Paumier, A., Onur, T.S., and Allen, N.J. (2023). Astrocyte contribution to dysfunction, risk and progression in neurodegenerative disorders. Nat Rev Neurosci 24, 23–39. 10.1038/s41583-022-00641-1.

8. Angel, J.P., and Daniels, B.P. (2022). Paradoxical roles for programmed cell death signaling during viral infection of the central nervous system. Curr Opin Neurobiol 77, 102629. 10.1016/j.conb.2022.102629.

9. Mangalmurti, A., and Lukens, J.R. (2022). How neurons die in Alzheimer’s disease: Implications for neuroinflammation. Curr Opin Neurobiol 75, 102575. 10.1016/j.conb.2022.102575.

10. Heckmann, B.L., Tummers, B., and Green, D.R. (2019). Crashing the computer: apoptosis vs. necroptosis in neuroinflammation. Cell Death Differ 26, 41–52. 10.1038/s41418-018-0195-3.

11. Morgan, M.J., and Kim, Y.S. (2022). Roles of RIPK3 in necroptosis, cell signaling, and disease. Exp Mol Med 54, 1695–1704. 10.1038/s12276-022-00868-z.

12. Daniels, B.P., Snyder, A.G., Olsen, T.M., Orozco, S., Oguin, T.H., 3rd, Tait, S.W.G., Martinez, J., Gale, M., Jr., Loo, Y.M., and Oberst, A. (2017). RIPK3 Restricts Viral Pathogenesis via Cell Death-Independent Neuroinflammation. Cell 169, 301-313 e311. 10.1016/j.cell.2017.03.011.

13. Daniels, B.P., Kofman, S.B., Smith, J.R., Norris, G.T., Snyder, A.G., Kolb, J.P., Gao, X., Locasale, J.W., Martinez, J., Gale, M., Jr., et al. (2019). The Nucleotide Sensor ZBP1 and Kinase RIPK3 Induce the Enzyme IRG1 to Promote an Antiviral Metabolic State in Neurons. Immunity 50, 64–76 e64. 10.1016/j.immuni.2018.11.017.

14. Chou, T.W., Chang, N.P., Krishnagiri, M., Patel, A.P., Lindman, M., Angel, J.P., Kung, P.L., Atkins, C., and Daniels, B.P. (2021). Fibrillar alpha-synuclein induces neurotoxic astrocyte activation via RIP kinase signaling and NF-kappaB. Cell Death Dis 12, 756. 10.1038/s41419-021-04049-0.

15. Wu, L., Chung, J.Y., Cao, T., Jin, G., Edmiston, W.J., 3rd, Hickman, S., Levy, E.S., Whalen, J.A., Abrams, E.S.L., Degterev, A., et al. (2021). Genetic inhibition of RIPK3 ameliorates functional outcome in controlled cortical impact independent of necroptosis. Cell Death Dis 12, 1064. 10.1038/s41419-021-04333-z.

16. Guo, H., Koehler, H.S., Mocarski, E.S., and Dix, R.D. (2022). RIPK3 and caspase 8 collaborate to limit herpes simplex encephalitis. PLoS Pathog 18, e1010857. 10.1371/journal.ppat.1010857.

17. Cervantes, P.W., Martorelli Di Genova, B., Erazo Flores, B.J., and Knoll, L.J. (2021). RIPK3 Facilitates Host Resistance to Oral Toxoplasma gondii Infection. Infect Immun 89. 10.1128/IAI.00021-21.

18. Preston, S.P., Allison, C.C., Schaefer, J., Clow, W., Bader, S.M., Collard, S., Forsyth, W.O., Clark, M.P., Garnham, A.L., Li-Wai-Suen, C.S.N., et al. (2023). A necroptosis-independent function of RIPK3 promotes immune dysfunction and prevents control of chronic LCMV infection. Cell Death Dis 14, 123. 10.1038/s41419-023-05635-0.

19. Gul-Hinc, S., Michno, A., Zysk, M., Szutowicz, A., Jankowska-Kulawy, A., and Ronowska, A. (2021). Protection of Cholinergic Neurons against Zinc Toxicity by Glial Cells in Thiamine-Deficient Media. Int J Mol Sci 22. 10.3390/ijms222413337.

20. Salmina, A.B. (2009). Neuron-glia interactions as therapeutic targets in neurodegeneration. J Alzheimers Dis 16, 485–502. 10.3233/JAD-2009-0988.

21. Ibrahim, A.M., Pottoo, F.H., Dahiya, E.S., Khan, F.A., and Kumar, J.B.S. (2020). Neuron-glia interactions: Molecular basis of alzheimer’s disease and applications of neuroproteomics. Eur J Neurosci 52, 2931–2943. 10.1111/ejn.14838.

22. Sheridan, G.K., and Murphy, K.J. (2013). Neuron-glia crosstalk in health and disease: fractalkine and CX3CR1 take centre stage. Open Biol 3, 130181. 10.1098/rsob.130181.

23. Roh, J.S., and Sohn, D.H. (2018). Damage-Associated Molecular Patterns in Inflammatory Diseases. Immune Netw 18, e27. 10.4110/in.2018.18.e27.

24. Gong, T., Liu, L., Jiang, W., and Zhou, R. (2020). DAMP-sensing receptors in sterile inflammation and inflammatory diseases. Nat Rev Immunol 20, 95–112. 10.1038/s41577-019-0215-7.

25. Venegas, C., and Heneka, M.T. (2017). Danger-associated molecular patterns in Alzheimer’s disease. J Leukoc Biol 101, 87–98. 10.1189/jlb.3MR0416-204R.

26. Sasaki, T., Liu, K., Agari, T., Yasuhara, T., Morimoto, J., Okazaki, M., Takeuchi, H., Toyoshima, A., Sasada, S., Shinko, A., et al. (2016). Anti-high mobility group box 1 antibody exerts neuroprotection in a rat model of Parkinson’s disease. Exp Neurol 275 Pt 1, 220–231. 10.1016/j.expneurol.2015.11.003.

27. Sathe, K., Maetzler, W., Lang, J.D., Mounsey, R.B., Fleckenstein, C., Martin, H.L., Schulte, C., Mustafa, S., Synofzik, M., Vukovic, Z., et al. (2012). S100B is increased in Parkinson’s disease and ablation protects against MPTP-induced toxicity through the RAGE and TNF-alpha pathway. Brain 135, 3336–3347. 10.1093/brain/aws250.

28. Kaur, J., Singh, H., and Naqvi, S. (2023). Intracellular DAMPs in Neurodegeneration and Their Role in Clinical Therapeutics. Mol Neurobiol 60, 3600–3616. 10.1007/s12035-023-03289-9.

29. Smeyne, R.J., and Jackson-Lewis, V. (2005). The MPTP model of Parkinson’s disease. Brain Res Mol Brain Res 134, 57–66. 10.1016/j.molbrainres.2004.09.017.

30. Munoz-Manchado, A.B., Villadiego, J., Romo-Madero, S., Suarez-Luna, N., Bermejo-Navas, A., Rodriguez-Gomez, J.A., Garrido-Gil, P., Labandeira-Garcia, J.L., Echevarria, M., Lopez-Barneo, J., and Toledo-Aral, J.J. (2016). Chronic and progressive Parkinson’s disease MPTP model in adult and aged mice. J Neurochem 136, 373–387. 10.1111/jnc.13409.

31. Meller, D., Eysel, U.T., and Schmidt-Kastner, R. (1994). Transient immunohistochemical labelling of rat retinal axons during Wallerian degeneration by a monoclonal antibody to neurofilaments. Brain Res 648, 162–166. 10.1016/0006-8993(94)91917-8.

32. Yandamuri, S.S., and Lane, T.E. (2016). Imaging Axonal Degeneration and Repair in Preclinical Animal Models of Multiple Sclerosis. Front Immunol 7, 189. 10.3389/fimmu.2016.00189.

33. Manivasagam, S., Williams, J.L., Vollmer, L.L., Bollman, B., Bartleson, J.M., Ai, S., Wu, G.F., and Klein, R.S. (2022). Targeting IFN-lambda Signaling Promotes Recovery from Central Nervous System Autoimmunity. J Immunol 208, 1341–1351. 10.4049/jimmunol.2101041.

34. Kim, S.T., Son, H.J., Choi, J.H., Ji, I.J., and Hwang, O. (2010). Vertical grid test and modified horizontal grid test are sensitive methods for evaluating motor dysfunctions in the MPTP mouse model of Parkinson’s disease. Brain Res 1306, 176–183. 10.1016/j.brainres.2009.09.103.

35. Zhang, W., Liu, J., Chen, Q., Ding, W., Li, S., and Ma, L. (2022). Identification of ADP/ATP Translocase 1 as a Novel Glycoprotein and Its Association with Parkinson’s Disease. Neurochem Res 47, 3355–3368. 10.1007/s11064-022-03688-9.

36. Liddelow, S.A., Guttenplan, K.A., Clarke, L.E., Bennett, F.C., Bohlen, C.J., Schirmer, L., Bennett, M.L., Munch, A.E., Chung, W.S., Peterson, T.C., et al. (2017). Neurotoxic reactive astrocytes are induced by activated microglia. Nature 541, 481–487. 10.1038/nature21029.

37. Yun, S.P., Kam, T.I., Panicker, N., Kim, S., Oh, Y., Park, J.S., Kwon, S.H., Park, Y.J., Karuppagounder, S.S., Park, H., et al. (2018). Block of A1 astrocyte conversion by microglia is neuroprotective in models of Parkinson’s disease. Nat Med 24, 931–938. 10.1038/s41591-018-0051-5.

38. Snyder, A.G., Hubbard, N.W., Messmer, M.N., Kofman, S.B., Hagan, C.E., Orozco, S.L., Chiang, K., Daniels, B.P., Baker, D., and Oberst, A. (2019). Intratumoral activation of the necroptotic pathway components RIPK1 and RIPK3 potentiates antitumor immunity. Sci Immunol 4. 10.1126/sciimmunol.aaw2004.

39. Orozco, S.L., Daniels, B.P., Yatim, N., Messmer, M.N., Quarato, G., Chen-Harris, H., Cullen, S.P., Snyder, A.G., Ralli-Jain, P., Frase, S., et al. (2019). RIPK3 Activation Leads to Cytokine Synthesis that Continues after Loss of Cell Membrane Integrity. Cell Rep 28, 2275–2287 e2275. 10.1016/j.celrep.2019.07.077.

40. Orozco, S., Yatim, N., Werner, M.R., Tran, H., Gunja, S.Y., Tait, S.W., Albert, M.L., Green, D.R., and Oberst, A. (2014). RIPK1 both positively and negatively regulates RIPK3 oligomerization and necroptosis. Cell Death Differ 21, 1511–1521. 10.1038/cdd.2014.76.

41. Lier, J., Streit, W.J., and Bechmann, I. (2021). Beyond Activation: Characterizing Microglial Functional Phenotypes. Cells 10. 10.3390/cells10092236.

42. Kenkhuis, B., Somarakis, A., Kleindouwel, L.R.T., van Roon-Mom, W.M.C., Hollt, T., and van der Weerd, L. (2022). Co-expression patterns of microglia markers Iba1, TMEM119 and P2RY12 in Alzheimer’s disease. Neurobiol Dis 167, 105684. 10.1016/j.nbd.2022.105684.

43. Xicoy, H., Wieringa, B., and Martens, G.J. (2017). The SH-SY5Y cell line in Parkinson’s disease research: a systematic review. Mol Neurodegener 12, 10. 10.1186/s13024-017-0149-0.

44. Nanini, H.F., Bernardazzi, C., Castro, F., and de Souza, H.S.P. (2018). Damage-associated molecular patterns in inflammatory bowel disease: From biomarkers to therapeutic targets. World J Gastroenterol 24, 4622–4634. 10.3748/wjg.v24.i41.4622.

45. Andersson, U., Wang, H., Palmblad, K., Aveberger, A.C., Bloom, O., Erlandsson-Harris, H., Janson, A., Kokkola, R., Zhang, M., Yang, H., and Tracey, K.J. (2000). High mobility group 1 protein (HMG-1) stimulates proinflammatory cytokine synthesis in human monocytes. J Exp Med 192, 565–570. 10.1084/jem.192.4.565.

46. Fischer, S., Nasyrov, E., Brosien, M., Preissner, K.T., Marti, H.H., and Kunze, R. (2021). Self-extracellular RNA promotes pro-inflammatory response of astrocytes to exogenous and endogenous danger signals. J Neuroinflammation 18, 252. 10.1186/s12974-021-02286-w.

47. Festoff, B.W., Sajja, R.K., van Dreden, P., and Cucullo, L. (2016). HMGB1 and thrombin mediate the blood-brain barrier dysfunction acting as biomarkers of neuroinflammation and progression to neurodegeneration in Alzheimer’s disease. J Neuroinflammation 13, 194. 10.1186/s12974-016-0670-z.

48. Banjara, M., and Ghosh, C. (2017). Sterile Neuroinflammation and Strategies for Therapeutic Intervention. Int J Inflam 2017, 8385961. 10.1155/2017/8385961.

49. Silvis, M.J.M., Kaffka Genaamd Dengler, S.E., Odille, C.A., Mishra, M., van der Kaaij, N.P., Doevendans, P.A., Sluijter, J.P.G., de Kleijn, D.P.V., de Jager, S.C.A., Bosch, L., and van Hout, G.P.J. (2020). Damage-Associated Molecular Patterns in Myocardial Infarction and Heart Transplantation: The Road to Translational Success. Front Immunol 11, 599511. 10.3389/fimmu.2020.599511.

50. Alisi, A., Carsetti, R., and Nobili, V. (2011). Pathogen- or damage-associated molecular patterns during nonalcoholic fatty liver disease development. Hepatology 54, 1500–1502. 10.1002/hep.24611.

51. Brajer-Luftmann, B., Nowicka, A., Kaczmarek, M., Wyrzykiewicz, M., Yasar, S., Piorunek, T., Sikora, J., and Batura-Gabryel, H. (2019). Damage-Associated Molecular Patterns and Myeloid-Derived Suppressor Cells in Bronchoalveolar Lavage Fluid in Chronic Obstructive Pulmonary Disease Patients. J Immunol Res 2019, 9708769. 10.1155/2019/9708769.

52. Tumburu, L., Ghosh-Choudhary, S., Seifuddin, F.T., Barbu, E.A., Yang, S., Ahmad, M.M., Wilkins, L.H.W., Tunc, I., Sivakumar, I., Nichols, J.S., et al. (2021). Circulating mitochondrial DNA is a proinflammatory DAMP in sickle cell disease. Blood 137, 3116–3126. 10.1182/blood.2020009063.

53. Santilli, F., Vazzana, N., Bucciarelli, L.G., and Davi, G. (2009). Soluble forms of RAGE in human diseases: clinical and therapeutical implications. Curr Med Chem 16, 940–952. 10.2174/092986709787581888.

54. Sokolove, J., and Lepus, C.M. (2013). Role of inflammation in the pathogenesis of osteoarthritis: latest findings and interpretations. Ther Adv Musculoskelet Dis 5, 77–94. 10.1177/1759720X12467868.

55. Goldstein, R.S., Gallowitsch-Puerta, M., Yang, L., Rosas-Ballina, M., Huston, J.M., Czura, C.J., Lee, D.C., Ward, M.F., Bruchfeld, A.N., Wang, H., et al. (2006). Elevated high-mobility group box 1 levels in patients with cerebral and myocardial ischemia. Shock 25, 571–574. 10.1097/01.shk.0000209540.99176.72.

56. Teismann, P., Sathe, K., Bierhaus, A., Leng, L., Martin, H.L., Bucala, R., Weigle, B., Nawroth, P.P., and Schulz, J.B. (2012). Receptor for advanced glycation endproducts (RAGE) deficiency protects against MPTP toxicity. Neurobiol Aging 33, 2478–2490. 10.1016/j.neurobiolaging.2011.12.006.

57. Kong, Z.H., Chen, X., Hua, H.P., Liang, L., and Liu, L.J. (2017). The Oral Pretreatment of Glycyrrhizin Prevents Surgery-Induced Cognitive Impairment in Aged Mice by Reducing Neuroinflammation and Alzheimer’s-Related Pathology via HMGB1 Inhibition. J Mol Neurosci 63, 385–395. 10.1007/s12031-017-0989-7.

58. Langeh, U., and Singh, S. (2021). Targeting S100B Protein as a Surrogate Biomarker and its Role in Various Neurological Disorders. Curr Neuropharmacol 19, 265–277. 10.2174/1570159X18666200729100427.

59. Brambilla, L., Martorana, F., Guidotti, G., and Rossi, D. (2018). Dysregulation of Astrocytic HMGB1 Signaling in Amyotrophic Lateral Sclerosis. Front Neurosci 12, 622. 10.3389/fnins.2018.00622.

60. Cristovao, J.S., Morris, V.K., Cardoso, I., Leal, S.S., Martinez, J., Botelho, H.M., Gobl, C., David, R., Kierdorf, K., Alemi, M., et al. (2018). The neuronal S100B protein is a calcium-tuned suppressor of amyloid-beta aggregation. Sci Adv 4, eaaq1702. 10.1126/sciadv.aaq1702.

61. Huttunen, H.J., Kuja-Panula, J., Sorci, G., Agneletti, A.L., Donato, R., and Rauvala, H. (2000). Coregulation of neurite outgrowth and cell survival by amphoterin and S100 proteins through receptor for advanced glycation end products (RAGE) activation. J Biol Chem 275, 40096–40105. 10.1074/jbc.M006993200.

62. Druse, M.J., Gillespie, R.A., Tajuddin, N.F., and Rich, M. (2007). S100B-mediated protection against the pro-apoptotic effects of ethanol on fetal rhombencephalic neurons. Brain Res 1150, 46–54. 10.1016/j.brainres.2007.02.092.

63. Businaro, R., Leone, S., Fabrizi, C., Sorci, G., Donato, R., Lauro, G.M., and Fumagalli, L. (2006). S100B protects LAN-5 neuroblastoma cells against Abeta amyloid-induced neurotoxicity via RAGE engagement at low doses but increases Abeta amyloid neurotoxicity at high doses. J Neurosci Res 83, 897–906. 10.1002/jnr.20785.

64. Sorci, G., Bianchi, R., Riuzzi, F., Tubaro, C., Arcuri, C., Giambanco, I., and Donato, R. (2010). S100B Protein, A Damage-Associated Molecular Pattern Protein in the Brain and Heart, and Beyond. Cardiovasc Psychiatry Neurol 2010. 10.1155/2010/656481.

65. Juranek, J., Mukherjee, K., Kordas, B., Zalecki, M., Korytko, A., Zglejc-Waszak, K., Szuszkiewicz, J., and Banach, M. (2022). Role of RAGE in the Pathogenesis of Neurological Disorders. Neurosci Bull 38, 1248–1262. 10.1007/s12264-022-00878-x.

66. Song, J., Lee, W.T., Park, K.A., and Lee, J.E. (2014). Receptor for advanced glycation end products (RAGE) and its ligands: focus on spinal cord injury. Int J Mol Sci 15, 13172–13191. 10.3390/ijms150813172.

67. Kim, J., Waldvogel, H.J., Faull, R.L., Curtis, M.A., and Nicholson, L.F. (2015). The RAGE receptor and its ligands are highly expressed in astrocytes in a grade-dependant manner in the striatum and subependymal layer in Huntington’s disease. J Neurochem 134, 927–942. 10.1111/jnc.13178.

68. Serrano, A., Donno, C., Giannetti, S., Peric, M., Andjus, P., D’Ambrosi, N., and Michetti, F. (2017). The Astrocytic S100B Protein with Its Receptor RAGE Is Aberrantly Expressed in SOD1(G93A) Models, and Its Inhibition Decreases the Expression of Proinflammatory Genes. Mediators Inflamm 2017, 1626204. 10.1155/2017/1626204.

69. Shi, J., Xu, H., Cavagnaro, M.J., Li, X., and Fang, J. (2021). Blocking HMGB1/RAGE Signaling by Berberine Alleviates A1 Astrocyte and Attenuates Sepsis-Associated Encephalopathy. Front Pharmacol 12, 760186. 10.3389/fphar.2021.760186.

70. Ponath, G., Schettler, C., Kaestner, F., Voigt, B., Wentker, D., Arolt, V., and Rothermundt, M. (2007). Autocrine S100B effects on astrocytes are mediated via RAGE. J Neuroimmunol 184, 214–222. 10.1016/j.jneuroim.2006.12.011.

71. Villarreal, A., Seoane, R., Gonzalez Torres, A., Rosciszewski, G., Angelo, M.F., Rossi, A., Barker, P.A., and Ramos, A.J. (2014). S100B protein activates a RAGE-dependent autocrine loop in astrocytes: implications for its role in the propagation of reactive gliosis. J Neurochem 131, 190–205. 10.1111/jnc.12790.

72. Ding, S., Wang, C., Wang, W., Yu, H., Chen, B., Liu, L., Zhang, M., and Lang, Y. (2021). Autocrine S100B in astrocytes promotes VEGF-dependent inflammation and oxidative stress and causes impaired neuroprotection. Cell Biol Toxicol. 10.1007/s10565-021-09674-1.

73. Przedborski, S., Jackson-Lewis, V., Naini, A.B., Jakowec, M., Petzinger, G., Miller, R., and Akram, M. (2001). The parkinsonian toxin 1-methyl-4-phenyl-1,2,3,6-tetrahydropyridine (MPTP): a technical review of its utility and safety. J Neurochem 76, 1265-1274. 10.1046/j.1471-4159.2001.00183.x.

74. Lin, Q.S., Chen, P., Wang, W.X., Lin, C.C., Zhou, Y., Yu, L.H., Lin, Y.X., Xu, Y.F., and Kang, D.Z. (2020). RIP1/RIP3/MLKL mediates dopaminergic neuron necroptosis in a mouse model of Parkinson disease. Lab Invest 100, 503–511. 10.1038/s41374-019-0319-5.

75. Dionisio, P.A., Oliveira, S.R., Gaspar, M.M., Gama, M.J., Castro-Caldas, M., Amaral, J.D., and Rodrigues, C.M.P. (2019). Ablation of RIP3 protects from dopaminergic neurodegeneration in experimental Parkinson’s disease. Cell Death Dis 10, 840. 10.1038/s41419-019-2078-z.

76. Wehn, A.C., Khalin, I., Duering, M., Hellal, F., Culmsee, C., Vandenabeele, P., Plesnila, N., and Terpolilli, N.A. (2021). RIPK1 or RIPK3 deletion prevents progressive neuronal cell death and improves memory function after traumatic brain injury. Acta Neuropathol Commun 9, 138. 10.1186/s40478-021-01236-0.

77. Ito, Y., Ofengeim, D., Najafov, A., Das, S., Saberi, S., Li, Y., Hitomi, J., Zhu, H., Chen, H., Mayo, L., et al. (2016). RIPK1 mediates axonal degeneration by promoting inflammation and necroptosis in ALS. Science 353, 603–608. 10.1126/science.aaf6803.

78. Faust, H., Lam, L.M., Hotz, M.J., Qing, D., and Mangalmurti, N.S. (2020). RAGE interacts with the necroptotic protein RIPK3 and mediates transfusion-induced danger signal release. Vox Sang 115, 729–734. 10.1111/vox.12946.

79. Boytard, L., Hadi, T., Silvestro, M., Qu, H., Kumpfbeck, A., Sleiman, R., Fils, K.H., Alebrahim, D., Boccalatte, F., Kugler, M., et al. (2020). Lung-derived HMGB1 is detrimental for vascular remodeling of metabolically imbalanced arterial macrophages. Nat Commun 11, 4311. 10.1038/s41467-020-18088-2.

80. Wen, S., Li, X., Ling, Y., Chen, S., Deng, Q., Yang, L., Li, Y., Shen, J., Qiu, Y., Zhan, Y., et al. (2020). HMGB1-associated necroptosis and Kupffer cells M1 polarization underlies remote liver injury induced by intestinal ischemia/reperfusion in rats. FASEB J 34, 4384–4402. 10.1096/fj.201900817R.

81. Minsart, C., Liefferinckx, C., Lemmers, A., Dressen, C., Quertinmont, E., Leclercq, I., Deviere, J., Moreau, R., and Gustot, T. (2020). New insights in acetaminophen toxicity: HMGB1 contributes by itself to amplify hepatocyte necrosis in vitro through the TLR4-TRIF-RIPK3 axis. Sci Rep 10, 5557. 10.1038/s41598-020-61270-1.

82. Meng, R., Gu, L., Lu, Y., Zhao, K., Wu, J., Wang, H., Han, J., Tang, Y., and Lu, B. (2019). High mobility group box 1 enables bacterial lipids to trigger receptor-interacting protein kinase 3 (RIPK3)-mediated necroptosis and apoptosis in mice. J Biol Chem 294, 8872–8884. 10.1074/jbc.RA118.007040.

83. Sparvero, L.J., Asafu-Adjei, D., Kang, R., Tang, D., Amin, N., Im, J., Rutledge, R., Lin, B., Amoscato, A.A., Zeh, H.J., and Lotze, M.T. (2009). RAGE (Receptor for Advanced Glycation Endproducts), RAGE ligands, and their role in cancer and inflammation. J Transl Med 7, 17. 10.1186/1479-5876-7-17.

84. Kierdorf, K., and Fritz, G. (2013). RAGE regulation and signaling in inflammation and beyond. J Leukoc Biol 94, 55–68. 10.1189/jlb.1012519.

85. Bianchi, R., Adami, C., Giambanco, I., and Donato, R. (2007). S100B binding to RAGE in microglia stimulates COX-2 expression. J Leukoc Biol 81, 108–118. 10.1189/jlb.0306198.

86. Lander, H.M., Tauras, J.M., Ogiste, J.S., Hori, O., Moss, R.A., and Schmidt, A.M. (1997). Activation of the receptor for advanced glycation end products triggers a p21(ras)-dependent mitogen-activated protein kinase pathway regulated by oxidant stress. J Biol Chem 272, 17810–17814. 10.1074/jbc.272.28.17810.

87. Lee, K.J., Yoo, J.W., Kim, Y.K., Choi, J.H., Ha, T.Y., and Gil, M. (2018). Advanced glycation end products promote triple negative breast cancer cells via ERK and NF-kappaB pathway. Biochem Biophys Res Commun 495, 2195–2201. 10.1016/j.bbrc.2017.11.182.

88. Marsche, G., Semlitsch, M., Hammer, A., Frank, S., Weigle, B., Demling, N., Schmidt, K., Windischhofer, W., Waeg, G., Sattler, W., and Malle, E. (2007). Hypochlorite-modified albumin colocalizes with RAGE in the artery wall and promotes MCP-1 expression via the RAGE-Erk1/2 MAP-kinase pathway. FASEB J 21, 1145–1152. 10.1096/fj.06-7439com.

89. Yatim, N., Jusforgues-Saklani, H., Orozco, S., Schulz, O., Barreira da Silva, R., Reis e Sousa, C., Green, D.R., Oberst, A., and Albert, M.L. (2015). RIPK1 and NF-kappaB signaling in dying cells determines cross-priming of CD8(+) T cells. Science 350, 328–334. 10.1126/science.aad0395.

90. Liu, J., Zhu, Z., Wang, L., Du, J., Zhang, B., Feng, X., and Zhang, G. (2020). Functional suppression of Ripk1 blocks the NF-kappaB signaling pathway and induces neuron autophagy after traumatic brain injury. Mol Cell Biochem 472, 105–114. 10.1007/s11010-020-03789-5.

91. Yu, P.W., Huang, B.C., Shen, M., Quast, J., Chan, E., Xu, X., Nolan, G.P., Payan, D.G., and Luo, Y. (1999). Identification of RIP3, a RIP-like kinase that activates apoptosis and NFkappaB. Curr Biol 9, 539–542. 10.1016/s0960-9822(99)80239-5.

92. Newton, K., Sun, X., and Dixit, V.M. (2004). Kinase RIP3 is dispensable for normal NF-kappa Bs, signaling by the B-cell and T-cell receptors, tumor necrosis factor receptor 1, and Toll-like receptors 2 and 4. Mol Cell Biol 24, 1464–1469. 10.1128/MCB.24.4.1464-1469.2004.

93. Murphy, J.M., Czabotar, P.E., Hildebrand, J.M., Lucet, I.S., Zhang, J.G., Alvarez-Diaz, S., Lewis, R., Lalaoui, N., Metcalf, D., Webb, A.I., et al. (2013). The pseudokinase MLKL mediates necroptosis via a molecular switch mechanism. Immunity 39, 443–453. 10.1016/j.immuni.2013.06.018.

94. Jackson-Lewis, V., and Przedborski, S. (2007). Protocol for the MPTP mouse model of Parkinson’s disease. Nat Protoc 2, 141–151. 10.1038/nprot.2006.342.

95. Daniels, B.P., Jujjavarapu, H., Durrant, D.M., Williams, J.L., Green, R.R., White, J.P., Lazear, H.M., Gale, M., Jr., Diamond, M.S., and Klein, R.S. (2017). Regional astrocyte IFN signaling restricts pathogenesis during neurotropic viral infection. J Clin Invest 127, 843–856. 10.1172/JCI88720.

