## Supplemental Material for "Neuronal DAMPs exacerbate neurodegeneration via astrocytic RIPK3 signaling"

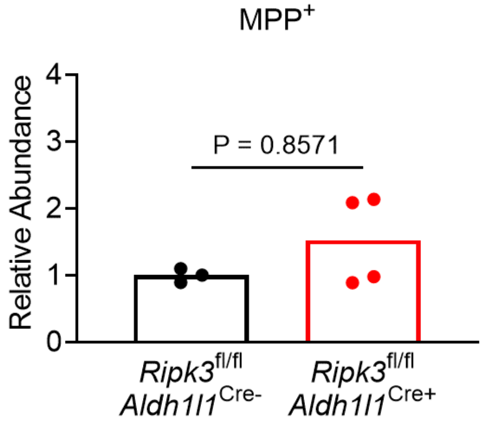


**Supplemental Figure 1.** ***Deletion of astrocytic Ripk3 does not impact MPTP metabolism in vivo.*** LC-MS analysis of $MPP^+$ abundance in midbrain homogenates of mice of indicated genotypes 90 minutes following intraperitoneal MPTP injection.

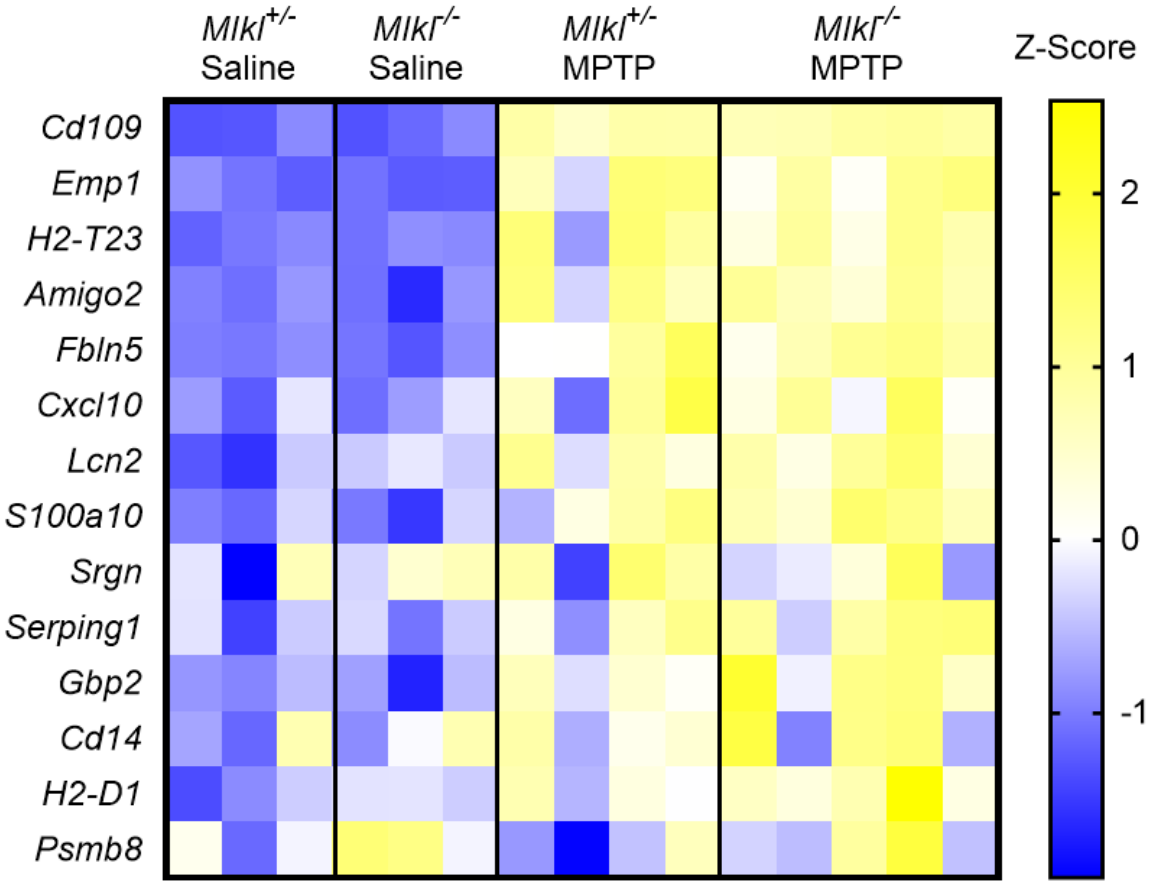


**Supplemental Figure 2.** ***Genes associated with astrocyte activation are not impacted by loss of MLKL in the MPTP model.*** qRT-PCR analysis of indicated genes in midbrain homogenates of *Mlkl* knockout and heterozygous littermate control animals 3 days post treatment with MPTP or saline.

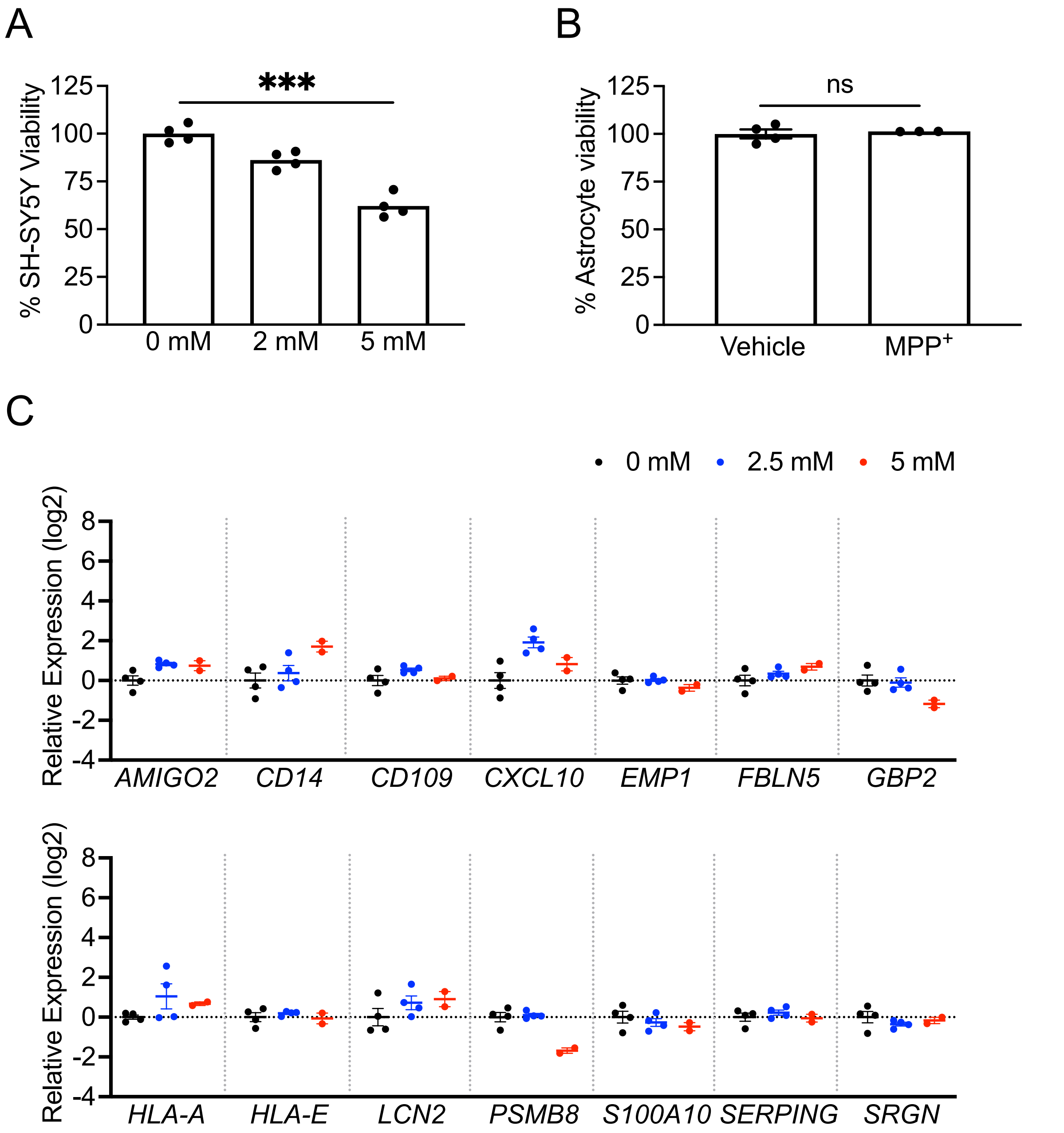


**Supplemental Figure 3.** ***MPP⁺ induces death in SH-SY5Y cells, but does not induce death or transcriptional activation in astrocytes.*** **(A-B)** Cell Titer Glo analysis of viability in SH-SY5Y **(A)** or primary human midbrain astrocyte **(B)** cultures treated with 2 or 5 mM **(A)** or 2.5 mM **(B)** $MPP^+$ for 24h. **(C)** qRT-PCR analysis of indicated genes in primary human midbrain astrocytes treated with 2.5 mM $MPP^+$ for 24h. *** $p<0.001$.

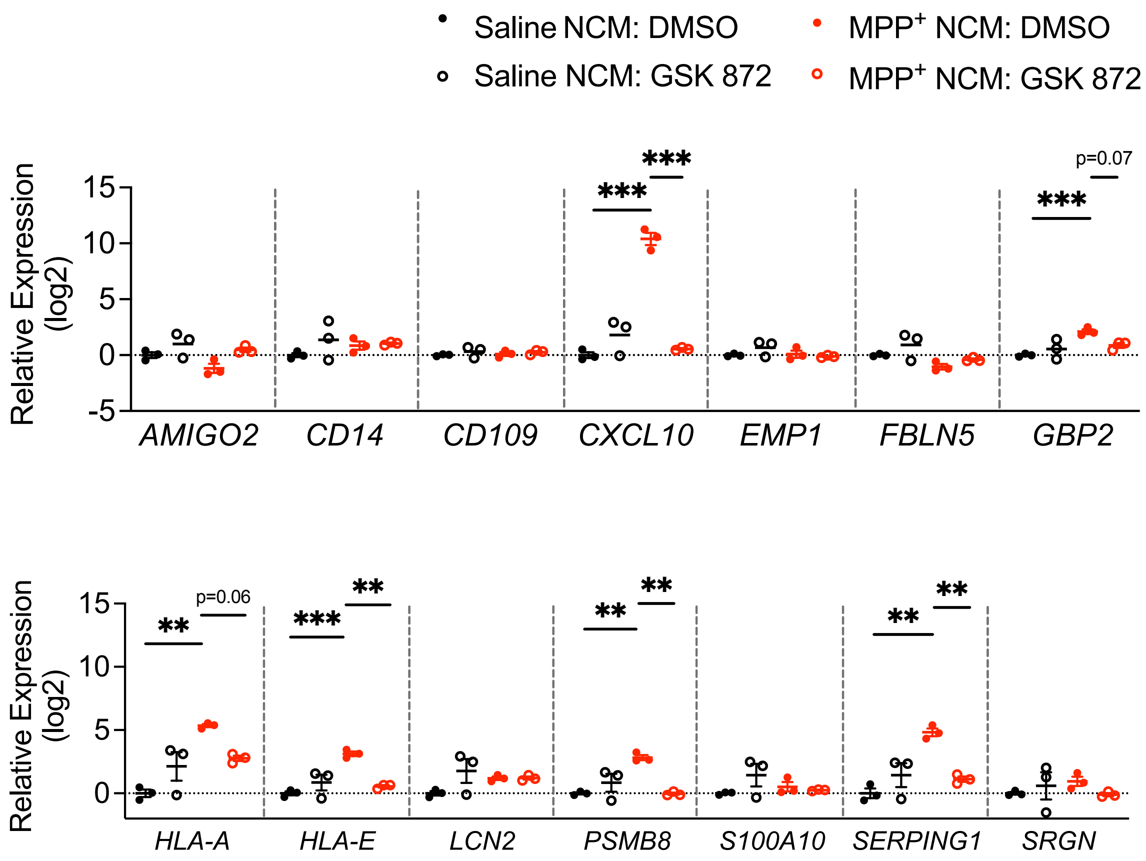


**Supplemental Figure 4.** ***Astrocytes maintain transcriptional activation for at least 24 hours following removal of MPP⁺ NCM.*** qRT-PCR analysis of indicated genes in primary human midbrain astrocytes treated for 24h with indicated conditions, followed by washing, addition of fresh culture medium (with no treatment), and an additional 24h incubation before harvest. *p<0.05, **p < 0.01, ***p < 0.001.

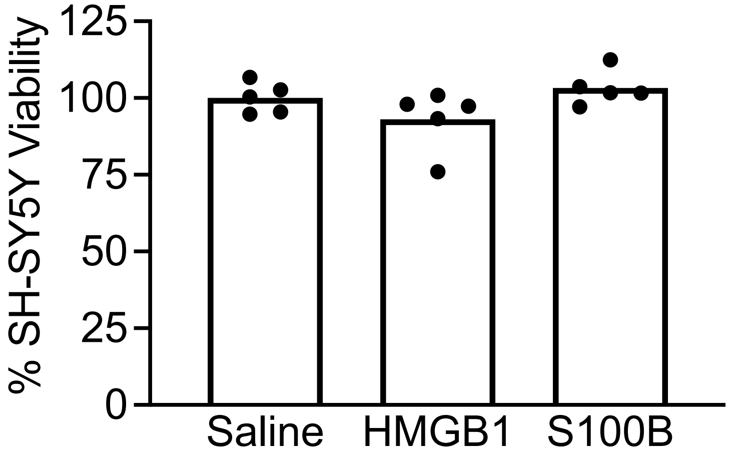


**Supplemental Figure 5.** ***Recombinant DAMPs are not intrinsically toxic to SH-SY5Y cells.*** **(A-B)** Cell Titer Glo analysis of viability in SH-SY5Y treated with indicated DAMP ligand for 24 hours.

### Supplementary Table S1.

#### Primer sequences for genotyping

| Target | Note | Primer Sequence (5'-3') | Product size |
|---|---|---|---|
| *Aldh1l1*-Cre/ERT2 | Internal control Forward | CTGTCCCTGTATGCCTCTGG | 415bp |
| | Internal control Reverse | AGATGGAGAAAGGACTAGGCTACA | |
| | Transgene Forward | CTTCAACAGGTGCCTTCCA | 198bp |
| | Transgene Reverse | GGCAAACGGACAGAAGCA | |
| *Mlkl*$^{-/-}$ | MLKL_001 | TATGACCATGGCAACTCACG | WT 498bp<br>KO 158bp |
| | MLKL_002 | ACCATCTCCCCAAACTGTGA | |
| | MLKL_003 | TCCTTCCAGCACCTCGTAAT | |
| *Ripk3*$^{-/-}$ | RIP3_001 | CGCTTTAGAAGCCTTCAGGTTGAC | WT 733bp<br>KO 485bp |
| | RIP3_002 | GCAGGCTCTGGTGACAAGATTCATGG | |
| | RIP3_003 | CCAGAGGCCACTTGTGTAGCG | |
| *Ripk3*$^{fl/fl}$ | R3FL_001 | ACGATGTCTTCTGTCAAGTTATG | WT 300bp<br>LoxP 334bp |
| | R3FL_002 | CAGTTCTTCACGGCTCAC | |
| | R3FL_003 | TCTGGTAAGGAGGGTCAC | |
| *Ripk3*-2xFV$^{fl/fl}$ | ROSA Forward | AGCACTTGCTCTCCCAAAGTC | 346bp |
| | ROSA Reverse | CCGACAAAACCGAAAATCTGTGGG | |
| | Transgene Forward | CGCTTTAGAAGCCTTCAGGTTGAC | 349bp |
| | Transgene Reverse | GCAGGCTCTGGTGACAAGATTCATGG | |

**Supplementary Table S2.**

**Primer sequences for qRT-PCR studies**

| Target | Forward | Reverse |
|---|---|---|
| *18S* hu | AGAAACGGCTACCACATCCA | CCCTCCAATGGATCCTCGTT |
| *18s* ms | CTTAGAGGGACAAGTGGCG | ACGCTGAGCCAGTCAGTGTA |
| *AMIGO2* hu | CTTCAGCGTTTGGAGGGCT | CAGGGAACAGTCACAGACAAAT |
| *Amigo2* ms | GAGGCGACCATAATGTCGTT | GCATCCAACAGTCCGATTCT |
| *CCL2* hu | GCAGCAAGTGTCCCAAAGAA | CTGGGGAAAGCTAGGGGAAA |
| *CD109* hu | CAGGAATGTGGACTCTGGGT | CTTTCGGACATGTGGACTGC |
| *CD109* ms | CACAGTCGGGAGCCCTAAAG | GCAGCGATTTCGATGTCCAC |
| *CD14* hu | CCGCTGTGTAGGAAAGAAGC | GCAGCGGAAATCTTCATCGT |
| *CD14* ms | GGACTGATCTCAGCCCTCTG | GCTTCAGCCCAGTGAAAGAC |
| *CXCL10* hu | GTGGCATTCAAGGAGTACCTC | TGATGGCCTTCGATTCTGGATT |
| *Cxcl10* ms | CCCACGTGTTGAGATCATTG | CACTGGGTAAAGGGGAGTGA |
| *EMP1* hu | CCAGTACACCAGCAGAGGAA | AACAGTAGCGATGTGGACCA |
| *Emp1* ms | GAGACACTGGCCAGAAAAGC | TAAAAGGCAAGGGAATGCAC |
| *FBLN5* hu | TCGCCAGTCAGGACAGTGT | AGTAGGGGTTCGAGTAGGGC |
| *Fbln5* ms | CTTCAGATGCAAGCAACAA | AGGCAGTGTCAGAGGCCTTA |
| *GBP2* hu | CTATCTGCAATTACGCAGCCT | TGTTCTGGCTTCTTGGGATGA |
| *Gbp2* ms | GGGGTCACTGTCTGACCACT | GGGAAACCTGGGATGAGATT |
| *HLA-A* hu | GACCAGGAGACACGGAATGTG | CCTCGTTCAAGGCGATGTAATC |
| *HLA-E* hu | TTCCGAGTGAATCTGCGGAC | GTCGTAGGCGAACTGTTCATAC |
| *H2-D1* ms | TCCGAGATTGTAAAGCGTGAAGA | ACAGGGCAGTGCAGGGATAG |
| *H2-T23* ms | GGACCGCGAATGACATAGC | GCACCTCAGGGTGACTTCAT |
| *LCN2* hu | GAAGTGTGACTACTGGATCAGGA | ACCACTCGGACGAGGTAACT |
| *Lcn2* ms | CCAGTTCGCCATGGTATTTT | CACACTCACCACCCATTCAG |
| *PSMB8* hu | GGTCCTACATTAGTGCCTTACGG | CGCAGATAGTACAGCCTGCATT |
| *Psmb8* ms | CAGTCCTGAAGAGGCCTACG | CACTTTCACCCAACCGTCTT |
| *S100A10* hu | ATGAAGGACCTGGACCAGTG | GCAGATTCCTTAAGCGACCC |
| *S100a10* ms | CCTCTGGCTGTGGACAAAAT | CTGCTCACAAGAAGCAGTGG |
| *SERPING1* hu | GGGATGCTTTGGTAGATTTCTCC | GAGGATGCTCTCCAGGTTTGT |
| *Serping1* ms | ACAGCCCCCTCTGAATTCTT | GGATGCTCTCCAAGTTGCTC |
| *SRGN* hu | GGACTACTCTGGATCAGGCTT | CAAGAGACCTAAGGTTGTCATGG |
| *Srgn* ms | GCAAGGTTATCCTGCTCGGA | TGGGAGGGCCGATGTTATTG |
